# Using Neutralization Landscapes to enumerate Antibody Behavior and Decompose Antibody Mixtures

**DOI:** 10.1101/2020.08.28.270561

**Authors:** Tal Einav, Adrian Creanga, Sarah F. Andrews, Adrian B. McDermott, Masaru Kanekiyo

**Author notes:** Co-first authors (Tal Einav, Adrian Creanga).

## Abstract

Antibodies constitute a key line of defense against the diverse pathogens we encounter in our lives. While the interactions between a single antibody and a single virus are routinely characterized in exquisite detail, the inherent tradeoffs between attributes such as potency and breadth remain unclear. In addition, there is a wide gap between the discrete interactions of single antibodies and how antibody mixtures collectively repel multiple viruses. Here, we develop a new form of antigenic cartography called a Neutralization Landscape that enumerates the full space of antibody-virus interactions for antibodies targeting the influenza hemagglutinin stem. This reference set of antibody behaviors transforms the potency-breadth tradeoff into a readily solvable geometry problem. Using the Neutralization Landscape, we decompose the collective neutralization from multiple antibodies to characterize the composition and functional properties of the stem antibodies within. Looking forward, this framework can leverage the serological assays routinely performed for influenza surveillance to analyze how an individual’s antibody repertoire evolves over time in response to vaccination or infection.

**One-Sentence Summary:** We describe the full range of behaviors for antibodies targeting the stem of influenza hemagglutinin, enabling us to decompose the collective behavior of antibody mixtures and characterize the individual antibodies within.

---

A key problem in immunology is to discover antibodies that can protect against a wide range of viruses. Yet it is difficult to quantify the inherent tradeoff between antibody potency (how well a virus is neutralized) and breadth (how many different viruses are neutralized). This tradeoff is especially important for rapidly-evolving viruses such as influenza, where we seek antibodies that are both highly-potent and broadly-neutralizing (*1–3*). Because these goals can be mutually exclusive, and because characterizing new antibodies is time- and resource-intensive, we need a framework that extrapolates the behavior of a few antibodies to describe all phenotypes.

The situation is further complicated in the context of multiple (polyclonal) antibodies, as in our immune system. With every infection or vaccination against the influenza virus, our antibody repertoire is reshaped, leading to a complex immune landscape whose ability to protect us from past and current strains is difficult to quantify (*4, 5*). While much effort has been devoted to measuring individual antibodies and predicting the effectiveness of their combinations (*6–9*), the inverse problem using the collective behavior of an antibody mixture to characterize the antibodies within is intractable without a framework to enumerate all antibody-virus interactions. Here, we create such a framework, opening up a unique perspective to describe the antibodies within mixtures.

To that end, we develop a new method to characterize antibody-virus interactions based on the techniques of antigenic cartography (*10, 11*) and antibody fingerprinting (*12*). Antigenic cartography creates a low dimensional map from hemagglutination inhibition titers that quantify how sera inhibit virus binding. While this technique imposes a structure for how sera can behave, it is unable to characterize the antibodies within a given serum nor predict the level of inhibition offered when sera are pooled together. Moreover, hemagglutination inhibition only characterizes antibodies binding to the head of influenza hemagglutinin (HA) and neglects antibodies targeting the HA stem, which generally inhibit a broader set of viruses (*1*) and which are being assessed in clinical trials as therapeutics (*13*).

A complementary method that partially offsets these drawbacks is antibody fingerprinting, which links the behavior of individual antibodies and antibody mixtures. The neutralization of large panels of antibodies are first clustered to identify patterns or “fingerprints” (*12*). By applying this process in reverse, neutralization from polyclonal sera can be decomposed to identify constituent antibodies from the original panel.

In this work, we create a Neutralization Landscape that characterizes the interaction between HA stem-targeting antibodies and influenza viruses. This approach pushes beyond cartography and fingerprinting in three key ways. First, as in cartography, we apply multidimensional scaling to antibody-virus measurements and project them into 2D, yet we do so at the level of individual antibodies rather than sera. By shifting the focus to antibodies, we quantify neutralization in absolute units without the need for normalization factors. Moreover, whereas the dominant antibodies within sera are generally unknown, and mapping polyclonal sera disrupts the map structure, the composition and binding site of individual antibodies can be precisely quantified. For these reasons, antibodies present a fundamental unit of the immune response that can more cleanly map virus interactions.

Second, this landscape serves as a discovery space for new antibodies and viruses. We empirically demonstrate that a 2D distance function (or metric) characterizes the >1000 antibody-virus interactions we measured. By positing that all other stem antibodies and viruses will be well characterized on the observed landscape, we can enumerate the range of possible behaviors. Thus, we can visualize the potency-breadth tradeoff and quantify how inhibiting more diverse viruses must decrease neutralization. In particular, we can predict the neutralization of the maximally potent antibody against any set of mapped viruses.

Finally, inspired by antibody fingerprinting, we develop a technique to decompose the collective neutralization from a mixture and characterize the antibodies within. Whereas previous efforts only detected specific patterns from a limited number of antibodies (*14*), our approach considers the full range of stem antibody behavior we extrapolate via the landscape. Using this rich set of behaviors, we determine the minimal number of stem antibodies (along with their full neutralization profile and stoichiometry) that could generate the observed signal from a mixture. Moreover, the Neutralization Landscape can remove the effects of non-HA-stem antibodies, and we validate such decompositions against 14 mixtures of HA head+stem antibodies. In this way, the Neutralization Landscape can peer into the influenza antibody response and quantify the stem antibodies within.

## Results

### Quantifying the Spectrum of Neutralization Profiles for Monoclonal Antibodies Targeting the Hemagglutinin Stem

Antigenic cartography utilizes metric multidimensional scaling to coalesce individual interactions (the ability of one antibody to inhibit one virus strain) into a global map (*15*). As a helpful geographic analogy, multidimensional scaling transforms pairwise distances between cities to create a state map (Figure S1). When cities are replaced by viruses and antibodies, the same procedure generates a map where the concentration of an antibody required to neutralize a virus is solely dictated by its distance to that virus, with smaller distances signifying more potent neutralization.

We assembled a virus panel comprising 24 H1N1 influenza strains collected between 1933-2018 and 27 H3N2 strains collected from 1968-2019 (Figure S2, Table S1). Neutralization was measured against 27 HA stem-targeting antibodies (Figure 1A, 17 previously published in (*16*) and 10 newly measured antibodies) representing major lineages of broadly neutralizing antibodies elicited by vaccination (*17–19*). The concentration of each antibody needed to neutralize every virus by 50% (the half maximal inhibitory concentration, IC_50_) was determined for most antibody-virus pairs (1136/1377=80%). Using these measurements, we constructed a neutralization landscape where the relative distance between an antibody and virus dictates the antibody’s potency (Figure 1B). As more viruses and antibodies are added, they lock into a fixed configuration, aside from global translations, rotations, and reflections.

**Figure 1.**
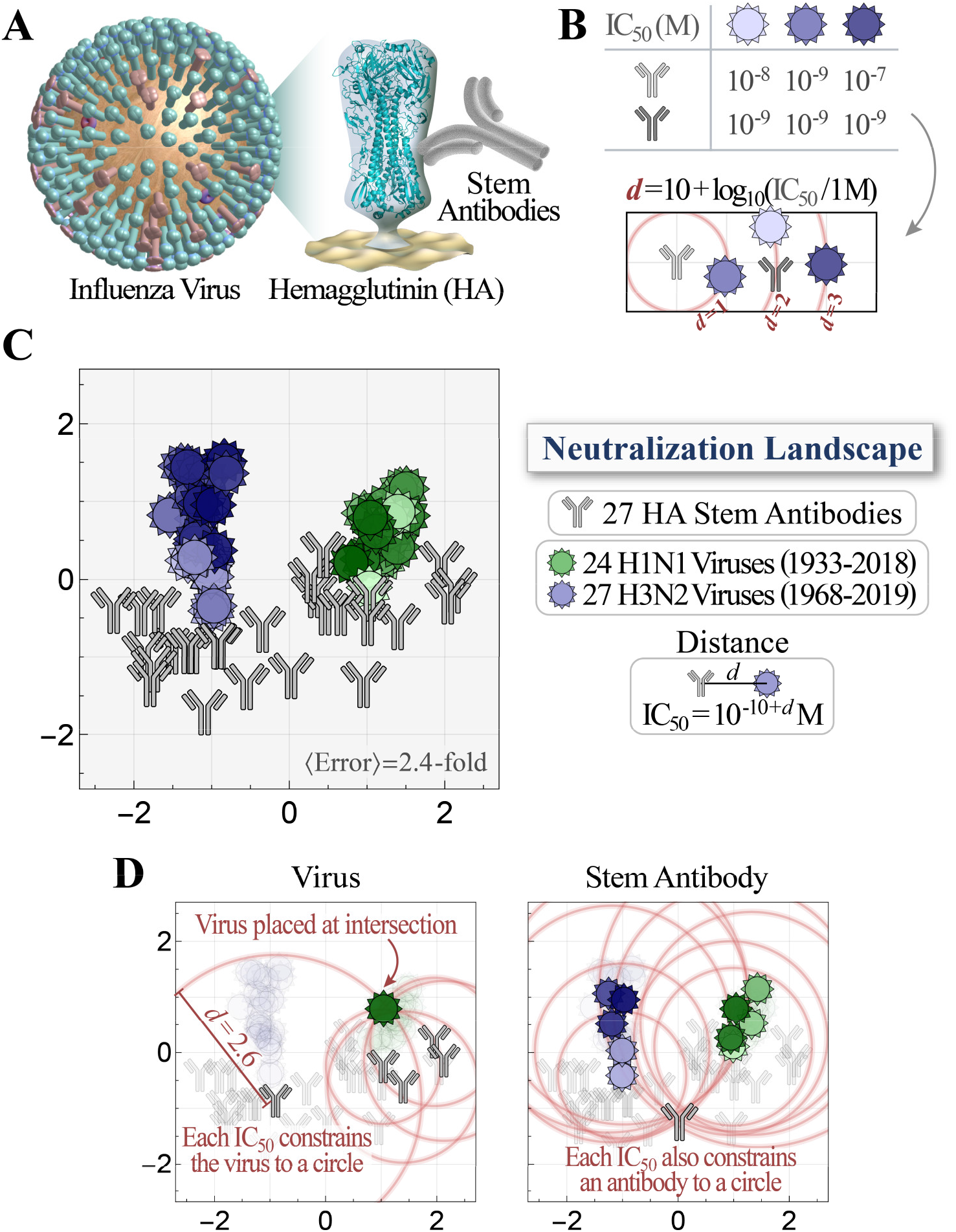
Neutralization Landscape for the influenza hemagglutinin stem. (A) Virus neutralization was measured for antibodies targeting the influenza HA stem. (B) Example showing how neutralization is transformed into a distance *d* on the landscape. An antibody with greater neutralization (*i.e.,* smaller half maximal inhibitory concentration, IC_50_) against a virus is placed closer to that virus. The light-gray antibody is positioned 1, 2, or 3 units away from the viruses, whereas the dark-gray antibody is 1 unit away from all three viruses. (C) Neutralization landscape of the HA stem quantifying the interactions between monoclonal antibodies (gray) and viruses (hues of green/blue; darker hues represent more recent viruses) (Figure S2, Table S1). A distance *d* between each antibody-virus pair corresponds to a neutralization IC_50_=10^-10+*d*^ M. Average error represents the mean fold-difference between the landscape IC_50_ values and measurements. (D) Examples of how a virus or antibody is positioned. Red circles represent the expected distance *d*=10+log_10_(IC_50_/1M), and an antibody or virus must lie as close as possible to the intersection of all circles.

Using these antibody-virus interactions, we created a Neutralization Landscape for the HA stem, with H1N1 and H3N2 viruses colored from lightest-to-darkest hues (oldest to more recent strains, Figure 1C). A distance *d* between an antibody and virus translates into an IC_50_ of 10^-10+*d*^ Molar (with 1 μg/mL = 6.6·10^-9^ M for the IgG antibodies considered here), so that greater distance represents exponentially decreasing inhibitory action. We quantified the error of the antibody and virus coordinates by computing the fold-error between the landscape IC_50_s and measured IC_50_s for all antibody-virus pairs, with a lower limit of 1-fold error for a landscape that perfectly represents the data (Methods). The 2D stem landscape had an 〈error〉=2.4-fold, comparable to the ≈2-fold accuracy of the neutralization assay. Surprisingly, when we remade the landscape in different dimensions, the error only decreased by 10% in 3D, although it more than tripled in 1D (Figure S3). Hence, we opt to represent the data in 2D which offers a balance between the ease of visualization and the accuracy of the landscape.

The resulting 2D landscape is described by 2·(27 antibodies + 51 viruses)=156 coordinates representing the 1136 antibody-virus measurements (156/1136=15% compression). To visualize the structure of the data that enables this compression, we draw circles of radius *d*=10+log_10_(IC_50_/1 Molar) around several antibodies measured against a virus. This virus must lie as close as possible to all circles, and its location can be determined via least squares minimization (Figure 1D, left panel). Antibodies and viruses are treated symmetrically, and hence an antibody is similarly fixed using its neutralization against multiple viruses (Figure 1D, right panel). We note that the circles shown in Figure 1D represent a small fraction of available data, with each virus constrained by 20 measurements and each antibody constrained by 40 measurements, on average. Error analysis shows that antibody and virus coordinates are tightly determined (Figure S2, Table S1). In summary, the Neutralization Landscape both visualizes and quantifies the interactions between our panel of antibodies and viruses, collapsing 1136 measurements spanning 3 orders of magnitude (from 6.8·10^-11^ M to 1.7·10^-7^ M).

### Triangulating New Antibodies or Viruses and Predicting their Behavior

The success of multidimensional scaling suggests that the interactions between stem antibodies and influenza viruses has a simple structure. If we postulate that this relationship holds between *all* stem antibodies and virus variants, then this landscape enumerates the full range of neutralization profiles. Every antibody or virus is represented by some point on the landscape, and its neutralization is defined by antibody-virus distance.

To test this hypothesis, we quantified how a new antibody or virus (whose measurements were not used to create the landscape) is represented on the map, and whether only a few of its measurements could predict its neutralization against the full virus panel. Thus, we withheld one antibody, reconstructed the Neutralization Landscape, triangulated the withheld antibody using 6 measurements, and compared its predicted versus measured neutralization against the remaining 45 viruses (Figure 2A, Methods). Notably, the reconstructed landscape looked nearly identical to the landscape using the full dataset (with all viruses shifting by <0.15 map units). Moreover, the subsequent triangulation correctly placed the antibody at the center of the landscape (only 0.15 map units from its location in Figure 1C).

**Figure 2.**
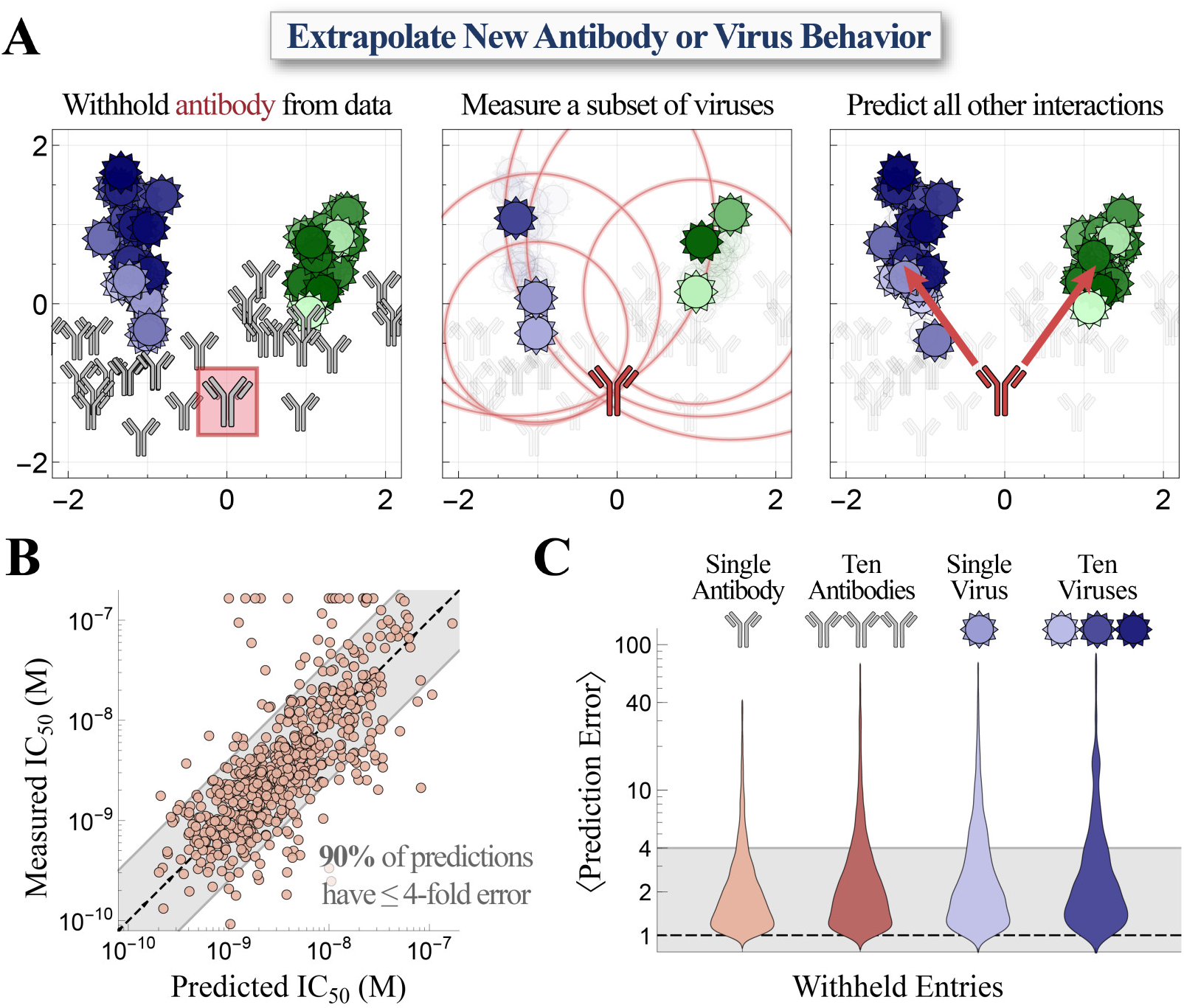
Extrapolating new antibody behavior. (A) *Left,* We withhold an antibody (CR8020, boxed in red) from our dataset and recreate the Neutralization Landscape. *Center,* The location of the withheld antibody is triangulated on the new landscape using a few measurements. *Right,* The neutralization against the remaining viruses is predicted using antibody-virus distance. (B) The 600 predicted versus measured IC_50_ values after withholding every antibody in our panel. Gray bands represent ≤ 4-fold error. (C) We withhold either antibodies or viruses (1 or 10 of each), triangulate each entity using a subset of measurements, and predict the remaining measurements. In each case, over 80% of predictions have ≤ 4-fold error.

We repeated this analysis, withholding each of the 27 antibodies in turn and predicting the left-out antibody’s complete neutralization profile. Collectively, 60% of the 600 predicted IC_50_s had ≤ 2-fold error, while 90% had ≤ 4-fold error (Figure 2B, left distribution in Figure 2C). When we similarly withheld and triangulated a virus, we found that 80% of the resulting predictions had ≤4-fold error (Figure 2C). As expected, prediction accuracy slowly improves when more measurements are used for triangulation (Figure S4).

As a more rigorous test, we next withheld multiple antibodies and viruses. We removed the 10 most recent viruses (isolated between 2010-2020), representing the practical scenario where past strains are used to infer the behavior of future variants. In addition, we assessed whether the landscape remained stable when antibodies from an entire region were depleted. Thus, we removed 10 antibodies from either the left half or right half of the map (removing more than a third of our 27 antibodies). In each scenario, we triangulated every entry as described above, using six measurements to predict the remaining data. We found that 80% of predictions had ≤ 4-fold error in all cases, demonstrating that the map can robustly infer new antibody or virus behavior (Figure 2C).

In summary, new antibodies or viruses can be added to the Neutralization Landscape using a few measurements. Once their positions are determined, we can quantify and visualize their response against all other mapped entries, and this predictive power increases as more antibodies and viruses are added to the landscape.

### Antibody-Virus Distance defines a Metric that Quantifies the Potency-Breadth Tradeoff

Although it is well-known that stem antibodies tend to neutralize a broader set of viruses than head antibodies (*1*, *20*), precisely quantifying the inherent tradeoff between antibody potency and breadth remains an open problem. Using the Neutralization Landscape, we transform this challenging biological question into a straightforward geometry problem.

A key mathematical property of the landscape is that antibody-virus distance forms a metric. In other words, our intuition for Euclidean geometry applies — for example, the antibody with the most potent neutralization (lowest IC_50_) against two viruses would lie exactly between them, minimizing the distance to either virus.

This setup is readily generalized to multiple viruses to answer a question that is intractable without a reference set for antibody behavior: How potently could any antibody neutralize *N* viruses? (Formally, what is the minimum 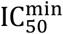 such that an antibody can exhibit an 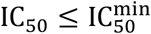 against all *N* viruses?) On the landscape, this optimal antibody lies at the center of the smallest circle bounding all *N* viruses.

To demonstrate this process, we considered the optimal antibody targeting all H1N1 or H3N2 vaccine strains from the 2004-05 to 2018-19 seasons (Methods). The optimal H3N2-specific antibody [blue] has a predicted 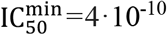 M, ~20x better than the best antibody in our panel [gray antibody 315-09-1B12] with a measured 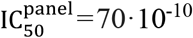 M (Figure 3A). In contrast, the optimal H1N1-specific antibody [green] has an 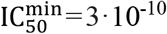 M, only 2.5x better than the 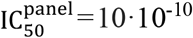 M from our best antibody [gray, antibody CR9114]. Hence, groups searching for better stem antibodies against these viruses should expect only marginal gains from additional H1N1-targeting antibodies, but far greater potential for finding new H3N2-targeting antibodies.

**Figure 3.**
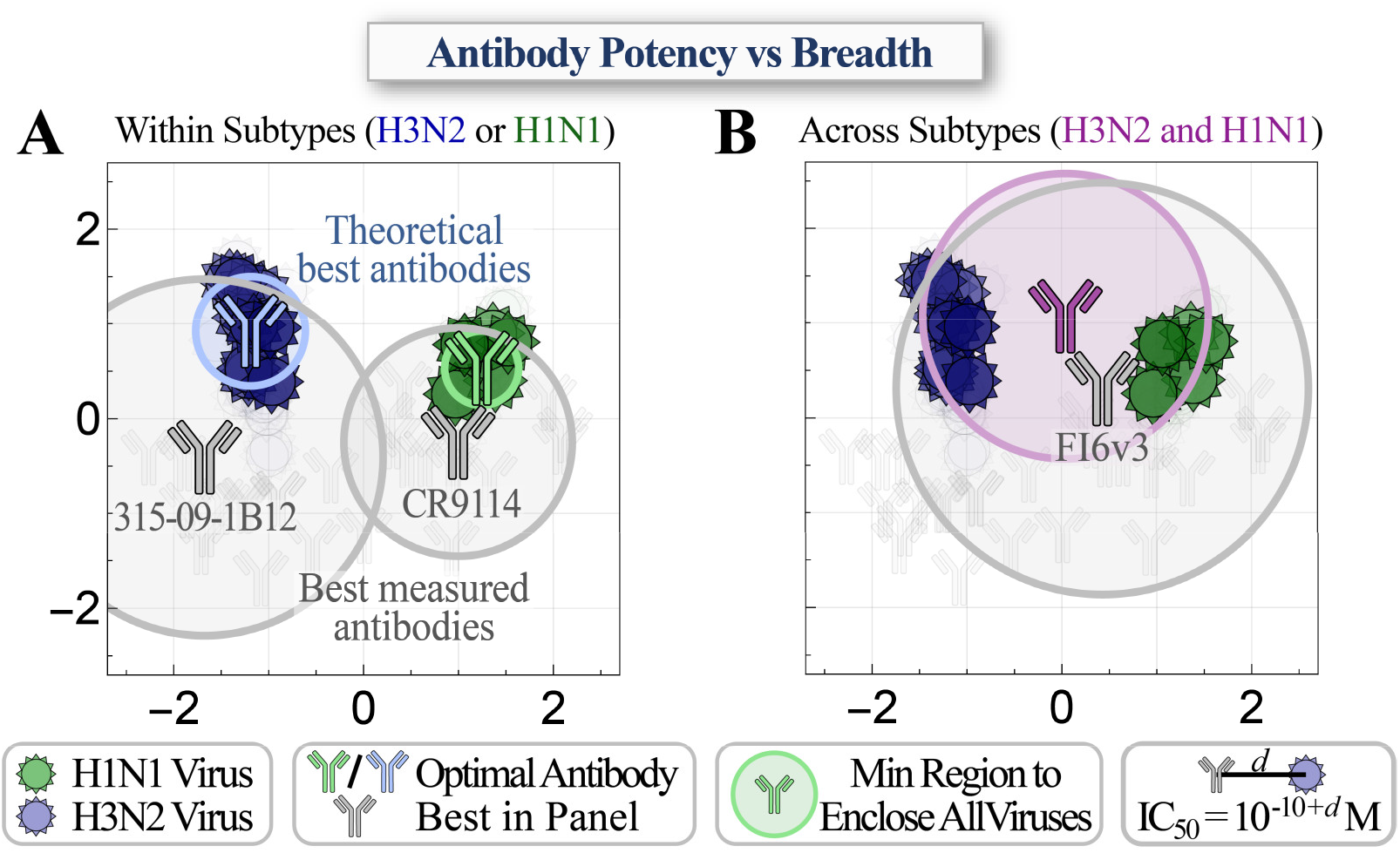
The neutralization profile of an optimal antibody against any set of mapped viruses. (A) Positions of the hypothetical highest-potency antibodies with the smallest IC_50_ against all H1N1 (green) or H3N2 (blue) vaccine strains from the 2004-05 to the 2018-19 seasons, compared to the best antibody in the panel (gray, labeled below with their names). A circle corresponding to the minimal IC_50_ enclosing all relevant viruses is drawn around each antibody. (B) Predicting the highest-potency antibody (purple) that neutralizes both the H1N1 and H3N2 strains compared to the best antibody in our panel (gray).

We can similarly assess how well a single antibody simultaneously neutralizes both the H1N1 and H3N2 vaccine strains. Because of the differences between these two subtypes, we expect that this enlarged breadth must come at the cost of decreased potency. Indeed, the Neutralization Landscape shows that a stem antibody can only exhibit an 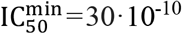 M against all of these viruses, ~5x worse that the best panel antibody with an 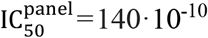 M [FI6v3] (Figure 3B).

This process is readily extended to any group of viruses, as well as to the more general question of how potently *N*_1_ antibodies could neutralize *N*_2_ viruses. Using the landscape, we can not only compute the optimal IC_50_, but also the specific neutralization profile against each virus on the panel. In this way, we can both visualize and quantify the full spectrum of antibody neutralization profiles and predict optimal antibody behavior.

As a technical note, a metric requires a triangle inequality. Since we only define the distance *d*_Ab-V_=10+log_10_(IC_50_/1 M) between an antibody (Ab) and virus (V), the usual triangle inequality becomes the quadrilateral inequality,

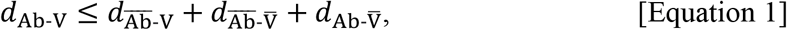

where 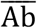 and 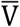 represent any other antibody or virus (Figure S5A). As with the traditional triangle inequality, this relationship codifies the notion that the distance between any Ab and V must be shorter than the next shortest route through 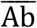 and 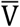. Altogether, there are 400,000 combinations of Ab, 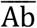, V, and 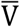 that can be directly tested against Equation 1 using the antibody-virus measurements (without requiring the Neutralization Landscape). We find that this inequality is satisfied in 99% of cases, demonstrating that our measurements are well described by the Euclidean metric. This empirical observation must be continually affirmed with future measurements.

### Isolating the Neutralization Profile of a Stem Antibody within a Head+Stem Mixture

By constraining the range of behaviors of a stem antibody, the Neutralization Landscape can also detect neutralization patterns from non-stem antibodies and remove those signals. For example, given the combined neutralization from an HA head+stem antibody mixture, we can predict the neutralization profile of the stem antibody alone (Figure 4A).

**Figure 4.**
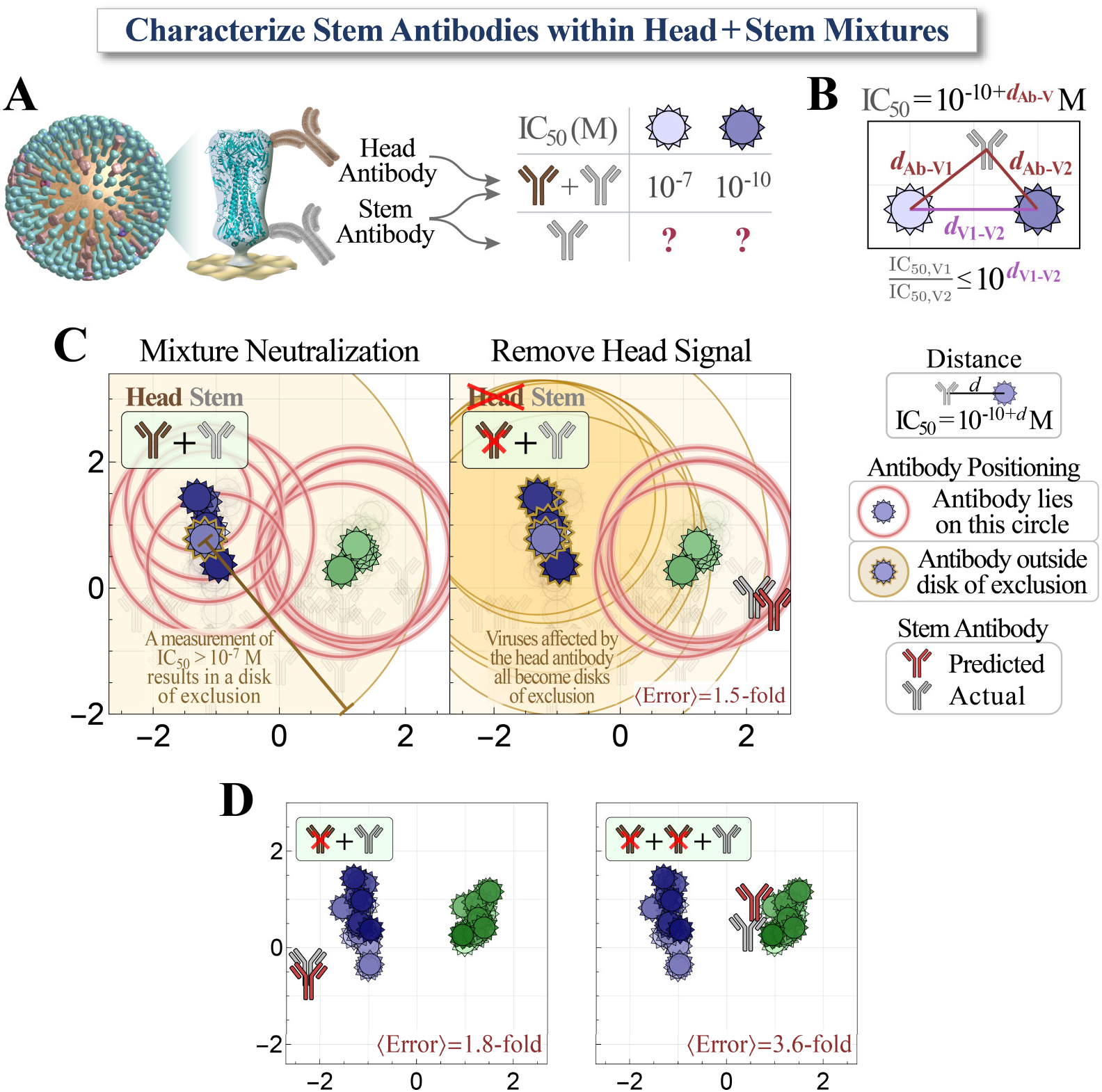
Characterizing a single stem antibody within head+stem mixtures. (A) We created mixtures of 2-3 antibodies targeting the HA head and stem. Using a mixture’s neutralization titers, we predict the behavior of the stem antibody within. (B) Virus-virus distance on the landscape constrains how differently any stem antibody can neutralize two viruses. (C) *Left,* The neutralization from a head+stem mixture (#7 in Figure S9) cannot be described by any point on the landscape due to the head antibody. The gold disk represents a lower bound on neutralization (*e.g.,* IC_50_ >10^-7^ M, *d*>3); an antibody should lie outside all gold disks but on every red circle (representing a defined IC_50_ value). *Right*, When the neutralization from the head antibody is removed, the IC_50_s of all blue viruses become lower bounds [gold disks], and we can infer the position of the stem antibody [red]. The insets at the top-left show the number of head [brown] and stem [gray] antibodies in each mixture, and whether the head signal has been removed [antibody icon crossed out]. (D) Examples showing additional mixtures combining 1 stem antibody with 1-2 head antibodies [additional decompositions in Figure S9]. Average error quantifies the fold-difference between the predicted antibody’s IC_50_s and measurements.

While antibody-virus distance on the landscape corresponds to neutralization, the distance between two viruses (V1 and V2) constrains how differently any stem antibody can neutralize both strains. In the extreme example where these viruses have identical HA stems and lie on the same coordinate (*d*_V1-V2_ = 0), every stem antibody will identically neutralize V1 and V2. Thus, if a head+stem antibody mixture neutralizes V1 far more potently than V2 (*e.g.*, with IC_50_s of 10^-10^ M and 10^-7^ M, respectively), this discrepancy must be caused by the head antibody increasing the mixture’s neutralization (decreases its IC_50_) against V1. The stem antibody alone should exhibit the same IC_50_ value ≥10^-7^ M against both viruses (a value larger than 10^-7^ M is possible if the head antibody increases the mixture’s neutralization against both viruses).

More generally, given a distance *d*_V1-V2_ between two viruses, any stem antibody (Ab) will obey |*d*_Ab-V1_ - *d*_Ab-V2_| ≤ *d*_V1-V2_ (Figure 4B) or equivalently

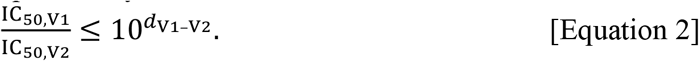

Given the combined neutralization from a head+stem mixture, we consider every pair of measurements (IC_50,V1_, IC_50,V2_) and determine whether their ratio exceeds 10^*d*_V_1_-V_2__^. If Equation 2 is satisfied, the measurements remain unchanged. Otherwise, the smaller value (say IC_50,V2_) is bounded below as per Equation 2 (IC_50,V2_ ≥ IC_50,V1_10^*d*_V_1_-V_2__^, Figure S5B-C; see Methods for how we account for noise in the measurements).

To test this process, we created 9 antibody mixtures (seven containing 1 head+1 stem antibody and two containing 2 head + 1 stem antibody) and measured them against our virus panel (*N*=321 data points, Figure S6). As an example, we consider one mixture where the stem antibody is H1N1-specific and does not neutralize any H3N2 viruses, whereas the head antibody neutralizes a few H1N1 and H3N2 strains (head antibody C05 + stem antibody CR6261). In combination, these two antibodies moderately neutralized some H3N2 viruses with an IC_50_ ≤ 10^-8^ M (Figure 4C left panel, red circles around the blue viruses), while others showed no detectable neutralization (virus in a gold disk represents an IC_50_ > 10^-7^ M outside our range of detection). The stem antibody should lie outside any gold disk while lying on the red circles representing IC_50_ values within our range of detection. As expected, no point on the landscape can satisfy these constraints in the left panel of Figure 4C, demonstrating that a stem antibody alone cannot give rise to the neutralization from this head+stem mixture.

As described above, we use the Neutralization Landscape to remove the effects of the head antibody. Given the close proximity of the H3N2 virus with no detectable neutralization (gold disk) to the H3N2 viruses with moderate neutralization (red circles), we correctly predict that the head antibody is responsible for this moderate neutralization. Thus, we increase the H3N2 IC_50_s and change them to lower bounds as dictated by Equation 2 (represented by gold disks in the right panel of Figure 4C). Notably, the H1N1 IC_50_s were unchanged by this process since the stem antibody’s neutralization dominated against these viruses, and hence the mixture’s H1N1 neutralization obeyed the constraints of the landscape.

With the head neutralization removed, we can triangulate the stem antibody on our map as discussed in the previous section (Figure 2A). In this way, we can characterize a stem antibody without knowing its individual neutralization profile nor the number or properties of the head antibodies in the combination. For our example mixture, the predicted stem antibody (red antibody in Figure 4C) lies near the true position of the stem antibody (gray), with an average 1.5-fold error between the predicted and measured IC_50_s across the virus panel. We repeated this analysis for our 9 antibody mixtures, combining either 1 or 2 head antibodies with a stem antibody, and found an 〈error〉 ranging between 1.5–5.2-fold (mean 3.4-fold, Figures 3D and S9). In summary, the set of stem antibody behaviors provided by the Neutralization Map can pick out the signature of a stem antibody within a mixture.

### Modeling and Characterizing Antibody Mixtures with Multiple Stem Antibodies

Given the possible behaviors for a single stem antibody, we predict how multiple stem antibodies act in concert, paving the way to explore a polyclonal antibody response (the purview of this section). In the reverse direction, we can use the collective neutralization from multiple stem antibodies to determine the number, stoichiometry, and neutralization profiles of the constituent antibodies (discussed in the following section).

To that end, we construct a biophysical model that calculates a mixture’s neutralization based on the neutralization of each individual antibody. Because the stem antibodies in our panel all target the same region of the HA stem (*17*, *21*, *22*), we treat their binding as competitive, where only one antibody can bind to an HA monomer at a time (Figure 5A). For two stem antibodies with individual neutralization 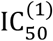 and 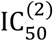, a mixture containing a fraction of the first antibody and *f*_2_ = 1 - *f*_1_ of the second antibody will have an

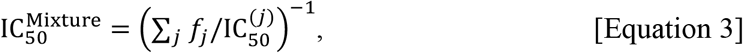

with this same equation holding for mixtures containing more antibodies (Methods).

**Figure 5.**
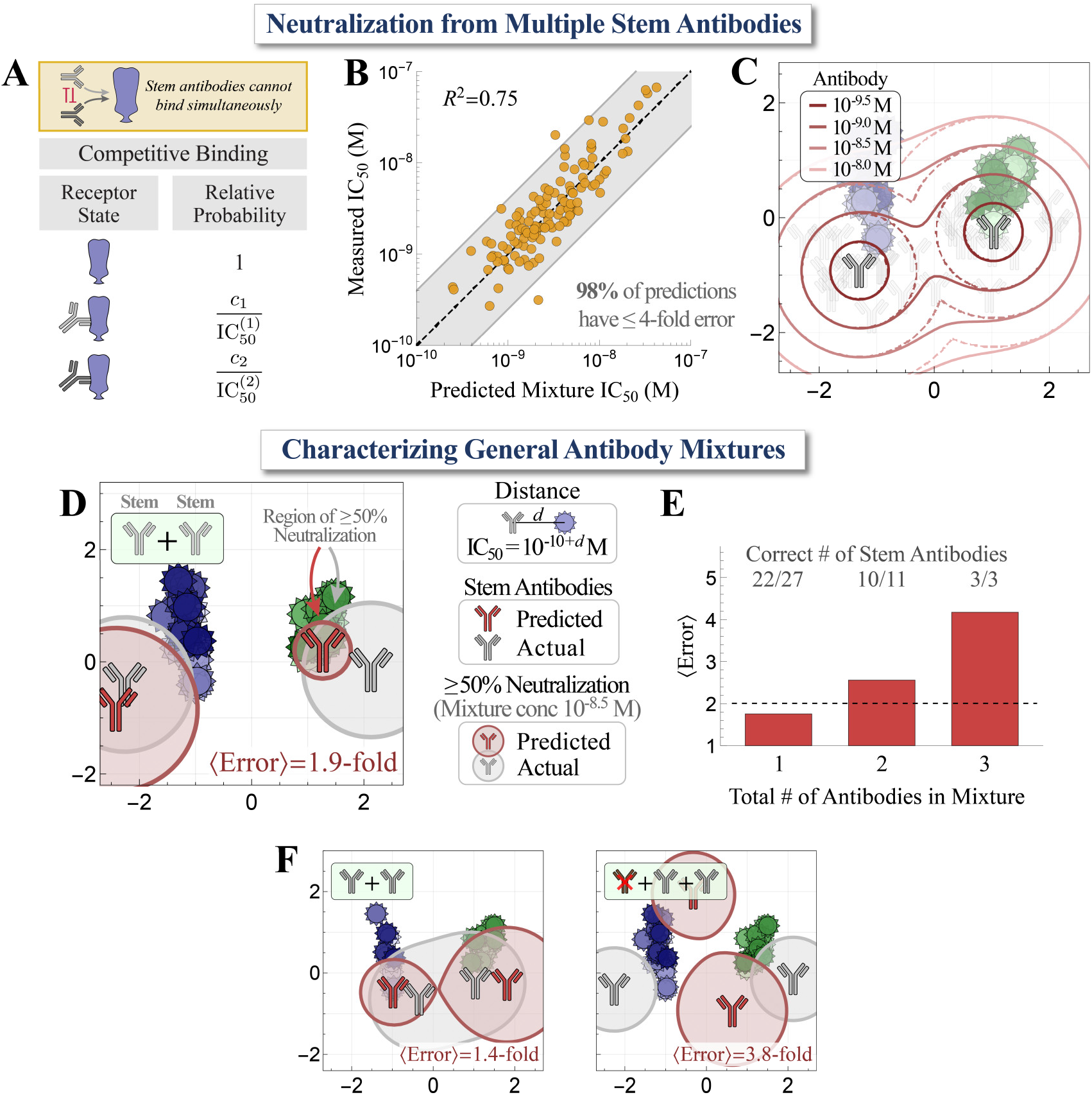
Characterizing mixtures with multiple stem antibodies. (A) Biophysical model for mixtures of stem antibodies binding competitively to an HA monomer. (B) We created stem+stem mixtures and measured them against the virus panel. We predict the mixture IC_50_ using the neutralization of each individual antibody. (C) Regions of ≥50% neutralization for a 2-antibody mixture [outlined in solid lines] versus the neutralization of each individual antibody [dashed lines]. The legend shows the total concentration of antibodies in the mixture. (D) Using a mixture’s neutralization titers, we predict the number, stoichiometry, and neutralization profiles of the stem antibodies within. One such stem+stem mixture is shown (gray antibodies, mixture #8 in Figure S9) together with the predicted decomposition (red). The circles around each antibody represent ≥50% virus neutralization when the total antibody concentration is 10^-8.5^ M while factoring in antibody stoichiometry within the mixture [Methods]. Average error represents the fold-difference between the collective neutralization predicted by the inferred stem antibodies and the measured neutralization of only the stem antibodies in the mixture. (E) Statistics for the decomposition of 27 monoclonal antibodies, 11 mixtures containing two antibodies (stem + stem or head+ stem), and 3 mixtures containing three antibodies (head+ head+ stem or head + stem+ stem). The fractions of decompositions that predicted the correct number of stem antibodies are shown above each bar. (F) Examples showing additional mixtures combining 2 stem antibodies with and without a head antibody [additional decompositions in Figure S9]. The regions around each antibody show the combined mixture neutralization, analogous to the solid lines in Panel C. The insets at the top-left show the number of head [brown] and stem [gray] antibodies in each mixture.

To test this competitive binding model, we created 4 stem+stem antibody mixtures and measured them against our virus panel (*N*=165 data points). Using the individual IC_50_s against each virus, we find tight agreement between the predicted and measured mixture IC_50_s, with 98% of predictions exhibiting ≤ 4-fold error (Figure 5B). On the landscape, such mixtures will neutralize a larger region [solid lines in Figure 5C] than either antibody alone [dashed lines]. In this way, we can use the competitive binding model to predict and visualize the behavior of general stem mixtures.

### Decomposing the Number and Functional Characteristics of Antibodies within Mixtures

By combining the results from the two previous sections, namely, removing the neutralization of head antibodies and enumerating the behavior of multiple stem antibodies, we can decompose general antibody mixtures with multiple head or stem antibodies. In essence, decomposition detects neutralization signatures that are impossible for a monoclonal antibody to achieve (*e.g.,* potent neutralization against viruses far apart on the landscape). In such cases, we search for the minimum number of antibodies that give rise to the apparent neutralization profile.

As an example, we measured the collective neutralization from a mixture of two stem antibodies against our virus panel (gray antibodies in Figure 5D; CR6261+ CR8020, 50%/50% composition). The algorithm scans through all possible configurations of *n*=1, 2, 3... antibodies on the landscape and determines which one best describes the mixture, terminating once the fold-error no longer appreciably decreases with additional antibodies (Figures S7-S8, Methods). This correctly predicted two stem antibodies (Figure 5D, red), although with a 10%/90% composition. The circle surrounding each antibody represents this fractional composition, so that the gray circles have the same radius whereas the red circle of the antibody on the left of the landscape (representing 90% composition) is larger than the red circle of the antibody on the right (10% composition). The areas covered by these circles represent ≥ 50% neutralization when the total mixture concentration equals a fixed amount we chose as 10^-8.5^ Molar. The deviation between the predicted and actual coordinates is partially compensated by predicting an uneven composition, so that the average fold-error between the measured and predicted IC_50_s against all viruses is 1.9-fold, comparable to experimental error. This demonstrates that the antibody response can be partially degenerate — where different mixtures exhibit similar behavior — since both the 50%/50% composition of gray antibodies and the 90%/10% composition of red antibodies exhibit similar neutralization.

We performed a similar analysis all our antibody mixtures, which included: (1) 27 monoclonal stem antibodies, (2) 11 mixtures containing two antibodies (4 stem+stem and 7 head+stem), and (3) 3 mixtures containing three antibodies (2 head+head+stem and 1 head+stem+stem) (Figure S6). As above, we blinded ourselves to each mixture’s composition, computationally removed any head antibody neutralization (even for mixtures containing only stem antibodies), determined which set of stem coordinates best characterized the neutralization profile, and compared the resulting neutralization against the true mixture compositions.

Across all mixtures, the collective neutralization from the stem antibodies was well characterized by these decompositions, with ≈2-fold error for the monoclonal antibodies and 3–4-fold error for the antibody mixtures (Figure 5E-F). For 22/27 monoclonal antibodies, the decomposition correctly predicted a single stem antibody, although 5 antibodies were overfit as 2-antibody mixtures due to deviations between landscape distances and measurements (Figure S9). All 7/7 head+stem mixtures and 3/4 stem+stem mixtures were correctly decomposed. The greatest deviations occurred either when a mixture contained two stem antibodies on the left and right edges of the landscape, resulting in a neutralization profile very similar to either a single stem antibody at the center of the landscape (2^nd^ panel from left of Figure S9B) or two stem antibodies along the central column (Figure 5F right panel, bottom panel of Figure S9C). Notably, even in cases where we inferred the incorrect number of stem antibodies, we found an 〈error〉 < 4-fold, implying that our predicted antibodies had a similar neutralization profile to the true antibodies. Collectively, these results suggest that while a mixture’s neutralization profile often uniquely identifies the stem antibodies within it (32/38=85% of cases), distinct mixtures can sometimes give rise to similar profiles, even when measured against a panel of 51 diverse viruses. The frequency of such degenerate mixtures, as well as the positions of additional viruses whose measurements would break these degeneracies, can be quantified by enumerating all possible mixtures on the landscape.

## Discussion

Since the advent of Fourier analysis in the 1800s, the ability to break signals into their simple underlying components has revolutionized diverse areas of science ranging from complex analysis to audio filtering to image reconstruction. In a recent example, live cells were imaged for the first time using six fluorescent probes whose emission spectra heavily overlapped, so that the net luminescence was a cacophony of signals (*23*). By characterizing each probe’s individual emission spectrum, the total signal could be unmixed, enabling six regions of live cells to be simultaneously viewed within a single cell. In the context of antibody-virus interactions, the challenge of unmixing lies both in enumerating the spectrum of antibody-virus behaviors as well as in decomposing the combined inhibition of antibody mixtures.

For the past two decades, antigenic maps of influenza have been created in order to visualize the interactions between sera and viruses, and it is common practice to determine each virus’s coordinates afresh for every dataset (*10*, *24*, *25*). Here, we take the opposite approach and create a Neutralization Landscape that fixes the positions of 27 HA stem antibodies and 51 influenza viruses, and then subsequently dissect the influenza antibody response using these positions. By constructing this Neutralization Landscape using monoclonal antibodies rather than sera, we gain a number of advantages. First, a monoclonal antibody is a precisely defined entity, whereas the composition of polyclonal sera is typically unknown. Thus, the structure of maps based on sera can be distorted by incorporating the effects of multiple antibodies, and such maps cannot explain asymmetries in the data (Figure S7, Methods). Second, serological assays for monoclonal antibodies quantify inhibitory *concentration* (in Molar or μg/mL units), so that map distance translates into these absolute units of measurement rather than to relative serum dilution units [Smith 2004]. On our landscape, distance imposes a metric on antibody-virus interactions.

As a result, the Neutralization Landscape serves as a discovery space for new antibodies, where a few measurements can pinpoint a new antibody or virus and predict its interaction with all other mapped entities (Figure 2). By systematically quantifying the range of behaviors for stem antibodies, this landscape can decompose polyclonal mixtures – even if they include antibodies binding to other epitopes such as the HA head – and quantify their fractional composition and neutralization profiles (Figures 3-4). Fundamentally, the information driving these predictions is in the *positions* of the viruses on the landscape, which quantifies the tradeoffs between antibody breadth and potency. When a mixture’s neutralization diverges from the possible profiles of a monoclonal antibody, it not only suggests that the mixture must be polyclonal but also presents a way to quantify the functional properties of the antibodies within the mixture.

With this approach, we can tackle one of the central problems in immunology, namely, using the collective neutralization from a mixture of antibodies to characterize the stem antibodies within. Although our mixtures only contained 1-3 HA head+stem antibodies, the human influenza antibody response is typically dominated by ≤ 5 antibodies (*26*) and in extreme cases by ≈1 antibody (*27*), suggesting that this level of resolution may be sufficient to decompose sera. This opens a number of applications including: (1) HA stem vaccines could be quantified in terms of both the fraction of elicited antibodies that target the HA stem as well as the neutralization profile of those antibodies (*28–32*). (2) Combining a neutralization landscape with a binding landscape (using antibody-virus dissociation constants) could quantify both neutralizing and nonneutralizing components of an antibody repertoire (*33*). (3) Given the inherently limited supply of each serum sample, we could rationally design the closest approximating antibody mixture using known antibodies, enabling broader studies of promising mixtures and facilitating the development of therapeutics.

This decomposition is inherently limited by the diversity of viruses used to probe a mixture. Our approach uses each virus as a “sensor” for nearby antibodies, so viruses should be widely spaced across the landscape to detect all possible antibodies. Due to experimental noise and the inaccuracy of the 2D representation, decomposition will only detect the dominant and distinct antibody signatures. An antibody comprising a small fraction of serum will minimally affect its neutralization and hence cannot be reliably detected. Moreover, antibodies with similar neutralization profiles may be represented by a single effective antibody. These cases add to degeneracy — a highly-understudied feature of the antibody response — where combinations of different antibodies give rise to similar functional responses.

An open question is whether the 2D Euclidean landscape presented here will suffice as more viruses and antibodies are added to the map, and if so, whether there is a biological reason why the potentially high-dimensional space of antibody-virus interactions is constrained to lie in a plane. We note that the positions of the viruses and antibodies exhibit distinct patterns on the landscape (Figure S2). H3N2 viruses roughly lie on a vertical line, with recent strains tending towards the top of the landscape. H1N1 viruses lie on a diagonal line with no clear temporal pattern. The stem antibodies lay on a horizontal strip, mostly at the bottom half of the landscape. These trends potentially indicate that the HA stem of H3N2 viruses is more tolerant to mutations than the HA stem of H1N1 viruses (*34*, *35*), and that the H3N2 stem may slowly mutate to evade immune pressure (*36*).

While the HA stem neutralizing antibodies used in this study all bind to the canonical stem super-epitope (*17*, *21*, *22*), antibodies targeting a new membrane-proximal stem epitope have recently been discovered (*37*). Future work can explore whether their behavior is captured by the existing landscape, or if a separate map is required for each epitope. It also remains to be seen whether there are portions of the landscape that antibodies or viruses cannot occupy (*e.g.,* because of viral protein stability or because an antibody would be self-reactive).

Looking forward, the analysis presented here serves as a stepping stone to track the stem antibody response over time and predict how our antibody repertoires will respond to a pathogen. How will the stem response evolve after multiple vaccinations or infections, and is there a path dependence to antibody development or is all the relevant information contained within the current antibody repertoire (*27*, *38–42*)? The Neutralization Landscape reframes this biological problem into a geometry problem, where antibody evolution can be studied as a dynamical system with perturbations imposed by vaccinations and infections. Taken together, this analysis serves as a new “computational microscope” to dissect our antibody repertoires and track how they evolve over the course of our lives.

## Supporting Information

The supporting information contains CSV files of: (1) the coordinates of the HA stem Neutralization Landscape, (2) neutralization measurements of the monoclonal antibodies and antibody mixtures, and (3) the full decompositions of all monoclonal antibodies and mixtures.

In addition, the supporting information contains a PDF file containing the aforementioned figures as well as a Mathematica notebook that contains the full analysis and reproduces all plots from this work.

## Acknowledgements

We would like to thank Jesse Bloom, Bernadeta Dadonaite, Alick Einav, Nathan Fridlyand, Vahe Galstyan, Ivelin Georgiev, Leslie Goo, Katie Gostic, Jane Kondev, Ivan Kosik, Lishibanya Mohapatra, Armita Nourmohammad, Jakub Otwinowski, Catherine Smith, Tyler Starr, and Jon Yewdell for their insights and input on this manuscript.

## Author contributions

Conceptualization and Methodology: T.E.; Investigation: T.E., A.C.; Writing – Review & Editing: T.E., A.C., S.A., M.K.

## Data and materials availability

All data is available in the supplementary materials, and the supplementary Mathematica notebook recreates all plots and analysis in this work.

## Supplementary Materials

Materials and Methods

Figures S1-S9

Table S1

ZIP folder containing all data

Mathematica notebook that creates the Neutralization Landscape and replicates all analysis

