## Supplementary Information for "Using Neutralization Landscapes to enumerate Antibody Behavior and Decompose Antibody Mixtures"


**Constructing the Neutralization Landscape** 2

- Applying Multidimensional Scaling to Create the Neutralization Landscape 2
- Applications and Limitations of the Neutralization Landscape 3
- Computing the Error of a Neutralization Landscape 5
- Utilizing Monoclonal Antibodies to Create the HA Stem Neutralization Landscape 5
- Patterns of Potential Virus Evolution based on Virus Positions 6
- Dimensionality of the Landscape 10
- Extrapolating the Behavior of New Antibodies 11
- The Triangle Inequality, Antibody-Virus Distance as a Metric 11
- Removing Head Antibody Neutralization 12
- The Tradeoff between Antibody Potency and Breadth 13

**Modeling Combinations of Antibodies** 15

- Overview of Antibody Mixtures 15
- The Collective Neutralization of Stem Antibody Mixtures is Well Characterized by Competitive Binding 16

**Decomposing Defined Antibody Mixtures** 18

- Why Polyclonal Mixtures need to be Represented using Multiple Points 19
- Defining Decomposition Error 20
- Overview of the Decomposition Algorithm 19
- Results from Decomposing Antibody Mixtures 21


The supporting information for this manuscript also includes the following CSV files:

- *(1) Neutralization Landscape Coordinates for the HA Stem.csv*
- *(2) Neutralization Data for Monoclonal Antibodies and Mixtures.csv*
- *(3) Decomposition Results.csv*

### A. Constructing the Neutralization Landscape

#### Applying Multidimensional Scaling to Create the Neutralization Landscape

To understand the utility of multidimensional scaling (MDS), we first introduce a useful analogy using a more familiar geographic map. Figure S1A shows a map of Texas along with several of its major cities. Using the scale bar at the top-right of this figure, it is straightforward to create a table of the distances between every pair of cities, as shown at the bottom of the panel.

Now consider the inverse scenario: given the distances between every pair of cities, how can you create a map of Texas? This is precisely the problem that multidimensional scaling solves. A simple algorithm is to choose random starting locations for each city and use numerical minimization techniques to minimize an error function such as $\sum_{j,k\in\text{cities}}(d_{jk}^{\text{Map}}-d_{jk}^{\text{Actual}})^2$ until the distance $d_{jk}^{\text{Map}}$ between cities $j$ and $k$ on the map matches their actual distance $d_{jk}^{\text{Actual}}$. This problem is also known as the sensor network localization problem, and it has been well studied in terms of how many distance measurements are required and whether the result is robust to noise (*1*).

Metric multidimensional scaling presumes there is an underlying structure for these distances. This is clear in our geographic analogy; for example, if a new city was drawn on the map, it could not simultaneously lie 10 km from Austin and 10 km from Houston, since those two cities are 150 km apart. The key insight from Smith *et al.* was that antibody-virus interactions may possess a similar underlying structure that could be exploited to create an analogous map (*2*). The ability of an antibody to inhibit a virus would be inversely proportional to its distance from that virus, with a smaller distance implying a more potent antibody. In this context, each row in the table represents an antibody, each column a virus, and each entry measures the inhibition of this antibody against the virus through a metric such as binding affinity, HAI, or neutralization.

Figure S1B shows an example that can be exactly mapped in 2D. The Euclidean distance $d$ between every antibody and virus (measured from the centers of their respective icons) corresponds to the 50% inhibitory concentration $IC_{50}$ (at which half of the virus is neutralized) of $10^{-10+d}$ Molar. A distance of $d=0$ between an antibody and virus represents an $IC_{50}\leq 10^{-10}$ M [which technically makes antibody-distance a pseudometric, since $d=0$ is not uniquely defined]. Since antibodies rarely bind with a dissociation constant less than $10^{-10}$ M (*3*, *4*), and because neutralization cannot occur without binding, there is a minimal loss in resolution from imposing this lower bound (which can be decreased further if needed). An antibody at a distance of $d=1$ from a virus exhibits an $IC_{50}=10^{-9}$ M, an antibody with distance $d=2$ exhibits an $IC_{50}=10^{-8}$ M, and so on, with a larger $IC_{50}$ indicating a less potent antibody.

Multidimensional scaling proceeds by minimizing the average error between the predicted and measured $IC_{50}$ values, $1/N\sum_{j\in\text{antibodies},k\in\text{viruses}}(d_{jk}^{\text{Predicted}}-d_{jk}^{\text{Measured}})^2$, where $N$ equals the number of measurements, $d_{jk}^{\text{Predicted}}$ represents the predicted map distance between antibody $j$ and virus $k$, and $d_{jk}^{\text{Measured}}=10+\log_{10}(IC_{50}^{(j,k)})$ denotes the experimentally measured $IC_{50}$ converted into map distance.

Note that the solution of multidimensional scaling will not be unique (since translations, rotations, and reflections do not affect antibody-virus distances), but beyond these rigid transformations, there can be different configurations of antibodies and viruses with similar error. As discussed in Smith *et al.*, local minima can be avoided by starting the minimization algorithm at multiple different starting configurations and choosing the resulting map with the smallest error (*2*).

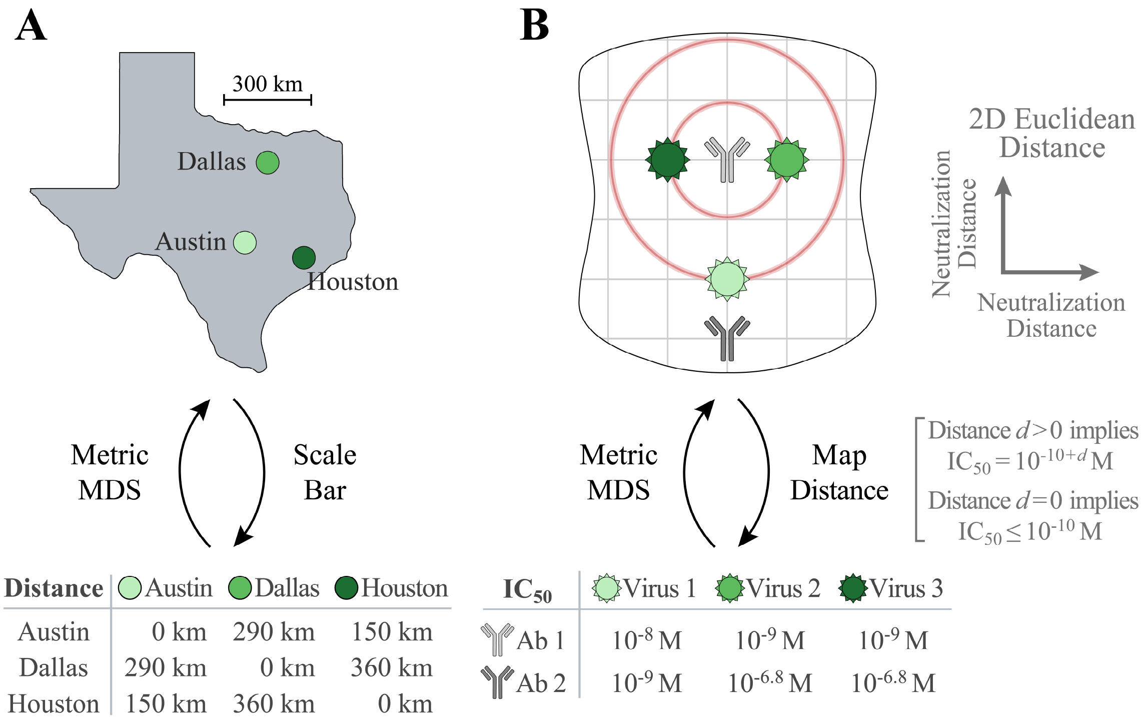


**Figure S1. Creating maps via multidimensional scaling.** (A) Metric multidimensional scaling (MDS) used in a geographic example to determine a map of Texas from a table of distances between cities. (B) In the context of antibody-virus interactions, each row of this table represents an antibody while each column represents a virus, with the entry of the table denoting the antibody's neutralization against the virus. Metric MDS determines coordinates for the antibodies and viruses, where a 2D Euclidean distance of 0 represents an $IC_{50}\leq 10^{-10}$ M while positive distance $d$ represents an $IC_{50}=10^{-10+d}$ M between any antibody-virus pair. Thus, the top antibody would neutralize all viruses within the smaller circle of radius 1 by at least 50% when at a concentration of $10^{-9}$ M, and it would neutralize all viruses within the larger circle of radius 2 by at least 50% when present at a concentration of $10^{-8}$ M.

### Applications and Limitations of the Neutralization Landscape

Given the widespread use of dimensionality reduction techniques (*e.g.*, multidimensional scaling, t-SNE, UMAP), it is worth emphasizing some applications of creating a framework that enumerates the full spectrum of antibody-virus responses (or some other biological context). These include:

- *Triangulating new antibodies/viruses*: Using ≈5 measurements for a new antibody or virus, we can fix its coordinate on the landscape and predict its full range of behavior (Figure 2). This approach can unify disparate datasets (*e.g.*, antibodies measured against viruses $V_1$-$V_5$ in one paper and $V_6$-$V_{10}$ in another paper) by placing all of them on the "level playing field" presented by a single Neutralization Landscape.
- *Design a replicate for a serum sample:* Given that the antibody repertoire is constantly changing, a serum sample is an inherently limited commodity. Unlike with monoclonal antibodies, there is currently no method to make more of a serum of interest. By characterizing the dominant antibody signatures within a serum, we can rationally design an approximation for this serum using known antibodies. More precisely, we can subtract the neutralization of head antibodies and

determine which stem antibodies on the map have the most similar neutralization profiles. Then, we can subtract the neutralization of those stem antibodies and determine which known head antibodies can give rise to the remaining signal. In this way, we can approximate all HA-targeting antibodies within a serum, and combine them with the appropriate stoichiometry to make the best possible approximation of the serum.

- *Antibody therapies and characterizing serum*: Given the increasing number of therapeutic antibodies under development (*5*, *6*), it is important to choose antibodies to cover as wide of a region on the landscape as possible with no holes and no redundancy. Similarly, groups seeking to understand serum responses should choose a diverse virus panel that covers a wide range of the map in order to detect as many distinct antibody signatures as possible.
- *Binding versus neutralization*: Although this work exclusively analyzes antibody neutralization, high-throughput assays can quantify antibody binding, providing complementary information on how well an antibody mixture binds to different viruses. It would be of great interest to compare landscapes created by binding versus neutralization measurements.
- *Degeneracy*: In this work, we found several instances where different antibody configurations can give rise to nearly identical neutralization profiles. This aspect of the immune response remains largely unexplored. How prevalent is this degeneracy, and how many different stem antibodies must be combined before the odds of correctly decomposing their collective response drops below 50% (see the fractions at the top of Figure 5E)?

In addition to these applications, we also mention some limitations of these dimensionality reduction approaches. First, dimensionality techniques characterize a given dataset, and it is not clear whether they can be extrapolated to characterize new behavior. While we demonstrate that a subset of our data can robustly predict the remainder of our measurements (Figure 2), additional viruses or antibodies may not conform to this same approach. Second, our assay only measured antibody neutralization, and hence remains ignorant of other mechanisms such as antibody-dependent cellular cytotoxicity or antibody-dependent cellular phagocytosis mediated through the Fc domain (*7*, *8*). Third, we only assessed H1N1 and H3N2 influenza viruses, and other subtypes could lie in vastly different regions of the landscape (or require a more complex metric or a higher dimension).

Finally, we emphasize one key difference between our approach (analyzing antibody-virus interactions) and traditional antigenic cartography (using serum-virus interactions), namely, that we do not assume any prior relationship between antibodies and viruses. Antigenic cartography assumes that serum elicited by infecting a ferret with virus *X* will be the maximally potent serum against virus *X* (more formally, that this serum is effectively a monoclonal antibody with the same coordinates as virus *X*). Thus, antigenic cartography cannot deal with asymmetries in inhibition data. For example, in the September 2020 WHO report (https://www.who.int/influenza/vaccines/virus/recommendations/202009_recommendation.pdf), serum raised against $V_1$=H1N1 A/California/07/2009 inhibited $V_2$=H1N1 A/Brisbane/02/2018 with a titer of 320, but serum raised against $V_2$ inhibited $V_1$ with a titer of 2560 (8-fold larger). Such asymmetries emphasize that these sera contain multiple antibodies, and that representing sera as single points on a map only approximates the full complexity of the system. In contrast, our approach allows both viruses and antibodies to reside anywhere on the map, thereby sidestepping this issue.

**Computing the Error of a Neutralization Landscape**

We applied multidimensional scaling to the monoclonal antibody neutralization data by minimizing the mean squared error between the predicted and measured $IC_{50}$ values,

$$\langle \text{map error}^2 \rangle = 1/N \sum_{j\in\text{antibodies},k\in\text{viruses}} (d_{jk}^{\text{Predicted}} - d_{jk}^{\text{Measured}})^2\, \Theta(d_{jk}^{\text{Predicted}}, d_{jk}^{\text{Measured}}),$$

using the number of measurements ($N$) and the predicted/measured antibody-virus map distances ($d_{jk}^{\text{Predicted}}/d_{jk}^{\text{Measured}}$) as described above. The function $\Theta(d_{jk}^{\text{Predicted}}, d_{jk}^{\text{Measured}})$ accounts for weak $IC_{50}$ values above the dynamic range of our assay ($d_{jk}^{\text{Measured}} > 1.6 \cdot 10^{-7}$ M, equivalent to 3.2 map units) so that the value of $d_{jk}^{\text{Predicted}}$ in such cases only contributes to the error when it falls below this bound. Following Smith *et al.*, we define $\Theta(d_{jk}^{\text{Predicted}}, d_{jk}^{\text{Measured}})=1$ when an exact antibody-virus $IC_{50}$ value is within the dynamic range of the assay ($d_{jk}^{\text{Measured}} \le 3.2$), whereas $\Theta(d_{jk}^{\text{Predicted}}, d_{jk}^{\text{Measured}}) = \frac{1}{1+e^{-10\left(d_{jk}^{\text{Predicted}} - d_{jk}^{\text{Measured}}\right)}}$ for $IC_{50}$s outside our dynamic range ($d_{jk}^{\text{Measured}}>3.2$) (*2*).

To relate this error function to the more intuitive $IC_{50}$ fold-error (equal to the ratio between the predicted and measured $IC_{50}$ values or its inverse, whichever is ≥1), we convert the square root of this mean squared error on the map into $IC_{50}$ ratios,

$$\langle \text{position error} \rangle \approx 10^{\sqrt{\langle \text{map error}^2 \rangle}}.$$

Note that mean-squared map error is the quantity minimized when creating the landscape, and the above relation demonstrates that minimizing the mean-squared map error will minimize the position error. However, once the MDS is complete, we compute the exact position error given by

$$\langle \text{position error} \rangle = 1/N \sum_{j\in\text{antibodies},k\in\text{viruses}} \text{fold-error}[IC_{50,jk}^{\text{predicted}}, IC_{50,jk}^{\text{measured}}]$$

where

$$\text{fold-error}[x,y] = \begin{cases} y/x, & x \le y \\ x/y, & x > y. \end{cases}$$

This is the error we show on our 2D landscape (gray text in the bottom-right of Figure 1C). ⟨Position error⟩=1-fold implies that the landscape perfectly represents the data, while 2-fold error implies that the landscape $IC_{50}$s will be between 2-fold larger and 2-fold smaller than the measured values, on average.

**Utilizing Monoclonal Antibodies to Create the HA Stem Neutralization Landscape**

Previous work by Creanga *et al.* measured the neutralization $IC_{50}$ of 18 HA stem-binding antibodies against 55 influenza strains (*9*). Replication-restricted reporter viruses (R3ΔPB1) were generated with the PB1 gene replaced by a fluorescent protein. Neutralization $IC_{50}$s were inferred by titrating the concentration of antibodies, with the $IC_{50}$ of each antibody-virus either fixed as the midpoint of the titration curve or bounded below as having $IC_{50}>1.6 \cdot 10^{-7}$ M (25μg/mL).

In this work, we focus exclusively on the H1N1 and H3N2 strains in the Creanga virus panel (49/55), to which we added the more recent strains H1N1 A/Idaho/07/2018 and H3N2 A/Perth/1008/2019. We measured an additional 10 stem antibodies and excluded one of the original stem antibodies (315-53-1A07) that did not neutralize any of the viruses. In total, the positions of these 51 viruses and 27 stem-binding antibodies are given in Table S1 as well as in the Supporting Information file named "*(1) Neutralization Landscape Coordinates - HA Stem.csv*".

In addition to quantifying the total error of the 2D landscape (discussed in the previous section), we compute the error of each entry (*i.e.*, each antibody or virus) by quantifying how far it can move before its error is increased by 2-fold. For each entry, we compute the Hessian of ⟨map error$^2$⟩ and find the direction $d_{max}$ of maximal increase (and the perpendicular direction $d_{min}$ where it minimally increases). We draw an ellipse whose semi-major axes ($r_{max}$ and $r_{min}$) in the maximal/minimal directions satisfied $\log_{10}(2)=½\, d_{max/min}\, r_{max/min}^2$, which corresponds to an ≈2-fold increase in map error. Across all antibodies and viruses, the uncertainty of all entries is always ≤0.4 units in any direction (Figure S2). We note that this uncertainty represents the *local* error of each antibody or virus about its map position, but there could be global minima elsewhere on the map where the entry would have similar error.

**Patterns of Potential Virus Evolution based on Virus Positions**
We color the H1N1 and H3N2 viruses on the Neutralization Landscape in hues from lightest (oldest) to darkest (more recent strains). Interestingly, the H3N2 viruses tend to travel upwards along the landscape, moving at 0.023 units/year on average (Figure S2B, with H3N2 A/Aichi/2/1968 the most notable exception). To put that into context, after 13 years, the antibodies below the H3N2 cluster would exhibit ≈2-fold less neutralization (since antibody-virus distance would increase by ≈0.3, and neutralization would decrease by $10^{0.3}$=2-fold). This HA stem evolution is markedly slower than the evolution of the highly-variable HA head (*10–12*), where hemagglutination inhibition decreases by approximately 2-fold each year (*2*, *13*).

Moreover, the HA stem of H1N1 viruses is nearly stationary, moving upwards by only 0.005 units/year on average for both the pre-pandemic and pandemic lineages (Figure S2C). At this pace, it would take 60 years for an antibody below the H1N1 cluster to exhibit 2-fold less neutralization. This discrepancy may be caused by the greater tolerance for HA stem mutations in H3N2 viruses (*14*, *15*). Moreover, H3N2 viruses acquire more mutations per year (*16*), and consequently, the H3N2 component of the influenza vaccine changed 8 times in the past decade, whereas the H1N1 component only changed 4 times.

The pattern in the positions of the H3N2 strains could represent virus evolution spurred by the HA-stem antibody response. While we note that broadly neutralizing anti-HA stem antibodies are rare in the human antibody repertoire, we note that the majority of these antibodies are positioned below the viruses (*17*). Moreover, the broadest neutralizing antibodies (*e.g.*, FI6v3, CR9114, MEDI8852, and 315-53-1B06) lie closer to the H1N1 viruses than to the H3N2 viruses consistent with previous findings that they evolved from unmutated common ancestors able to bind only group 1 HAs (*e.g.*, H1, H2, or H5) and acquired the ability to neutralize H3N2 viruses through somatic hypermutation.

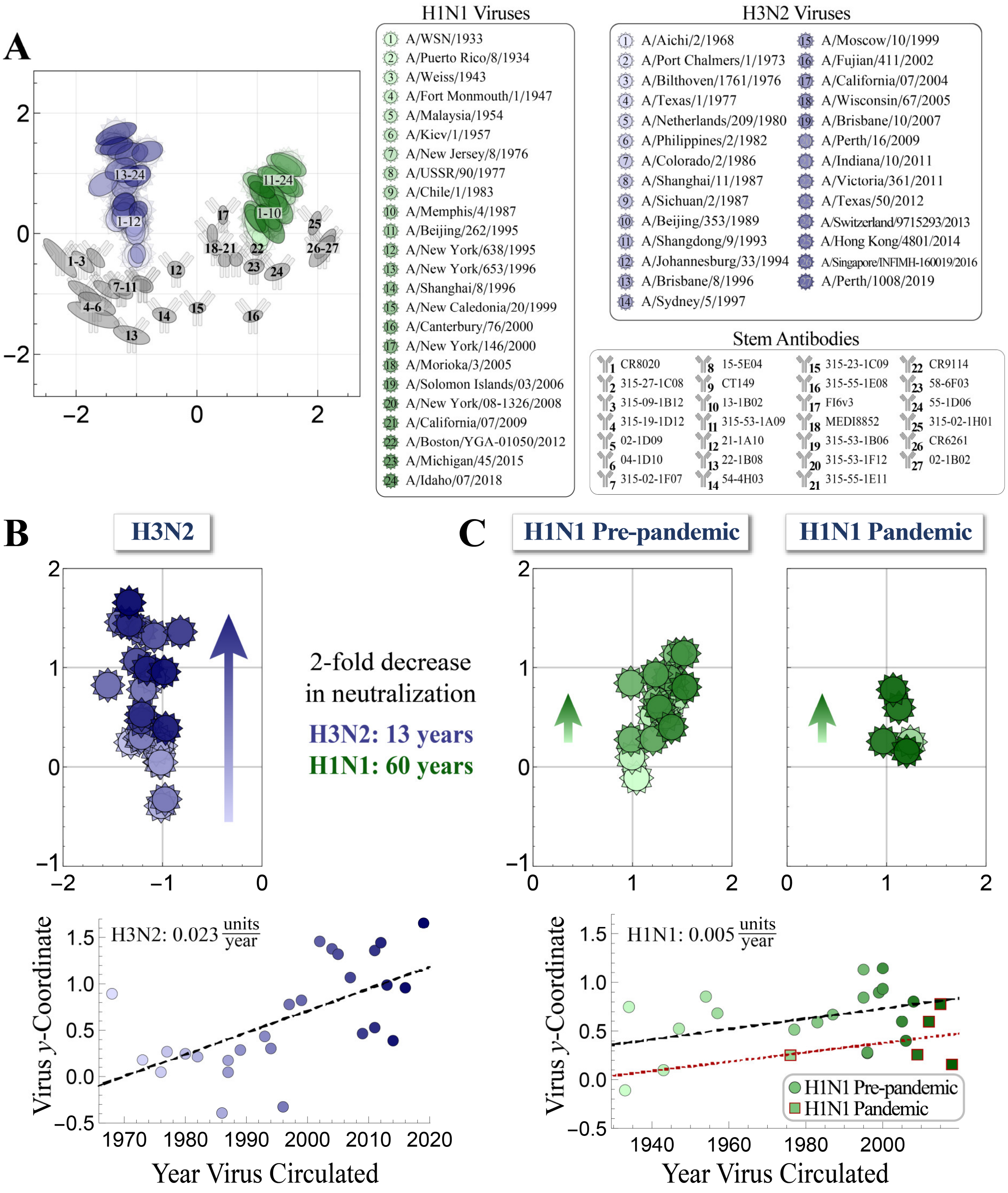


**Figure S2. Temporal patterns in virus neutralization.** (A) The coordinates of each antibody and virus, overlaid with an ellipse showing the error of each coordinate (denoting a 2-fold increase in the entry's error). The antibody numbering used in this figure is followed throughout this work in all figures and tables, so that Stem Ab 1 represents CR8020, Stem Ab 2 represents 315-27-1C08, and so on. (B-C) *Top*, Close up of the H3N2 or H1N1 viruses on the landscape, showing that more recent H3N2 strains lie towards the top of the landscape. For H1N1, we separate the pandemic lineage (A/New Jersey/8/1976 and all viruses from 2009 or later) from the pre-pandemic lineage. *Bottom*, *y*-coordinate for each virus versus its year of circulation, together with the best fit line (dashed black lines or the dotted red line for the pandemic H1N1 lineage).

**Table S1. Coordinates of the viruses and antibodies (Abs) on the Neutralization Landscape.** For each entry, we specify its map coordinates along with the size of the semi-major and semi-minor axis of the uncertainty ellipse shown in Figure S2A (subscript θ given in radians counterclockwise from the *x*-axis). The head antibodies used in some of the antibody mixtures are listed, although they are not positioned on the stem landscape.

| Virus or Antibody | Coordinate | Large Error (Angle) | Small Error (Angle) |
|---|---|---|---|
| H1N1 A/WSN/1933 | (1.04, -0.11) | 0.18 (θ=1.97) | 0.11 (θ=0.40) |
| H1N1 A/Puerto Rico/8/1934 | (1.30, 0.75) | 0.30 (θ=2.42) | 0.17 (θ=0.85) |
| H1N1 A/Weiss/1943 | (0.99, 0.10) | 0.33 (θ=2.02) | 0.16 (θ=0.45) |
| H1N1 A/Fort Monmouth/1/1947 | (1.20, 0.52) | 0.21 (θ=2.38) | 0.13 (θ=0.81) |
| H1N1 A/Malaysia/1954 | (1.40, 0.85) | 0.33 (θ=2.54) | 0.18 (θ=0.97) |
| H1N1 A/Kiev/1/1957 | (1.42, 0.68) | 0.25 (θ=2.42) | 0.14 (θ=0.85) |
| H1N1 A/New Jersey/8/1976 | (1.24, 0.25) | 0.24 (θ=2.11) | 0.13 (θ=0.54) |
| H1N1 A/USSR/90/1977 | (1.33, 0.51) | 0.24 (θ=2.27) | 0.13 (θ=0.70) |
| H1N1 A/Chile/1/1983 | (1.29, 0.59) | 0.23 (θ=2.34) | 0.14 (θ=0.77) |
| H1N1 A/Memphis/4/1987 | (1.24, 0.67) | 0.23 (θ=2.43) | 0.13 (θ=0.86) |
| H1N1 A/Beijing/262/1995 | (1.44, 1.13) | 0.31 (θ=2.69) | 0.15 (θ=1.12) |
| H1N1 A/New York/638/1995 | (0.98, 0.84) | 0.25 (θ=2.58) | 0.14 (θ=1.01) |
| H1N1 A/New York/653/1996 | (0.98, 0.28) | 0.30 (θ=2.13) | 0.16 (θ=0.56) |
| H1N1 A/Shanghai/8/1996 | (1.20, 0.28) | 0.24 (θ=2.15) | 0.13 (θ=0.58) |
| H1N1 A/New Caledonia/20/1999 | (1.36, 0.90) | 0.33 (θ=2.56) | 0.18 (θ=0.99) |
| H1N1 A/Canterbury/76/2000 | (1.23, 0.93) | 0.25 (θ=2.59) | 0.14 (θ=1.02) |
| H1N1 A/New York/146/2000 | (1.52, 1.14) | 0.32 (θ=2.67) | 0.15 (θ=1.10) |
| H1N1 A/Morioka/3/2005 | (1.27, 0.60) | 0.29 (θ=2.27) | 0.17 (θ=0.70) |
| H1N1 A/Solomon Islands/03/2006 | (1.39, 0.40) | 0.27 (θ=2.14) | 0.16 (θ=0.57) |
| H1N1 A/New York/08-1326/2008 | (1.52, 0.80) | 0.27 (θ=2.51) | 0.15 (θ=0.94) |
| H1N1 A/California/07/2009 | (0.97, 0.26) | 0.26 (θ=2.13) | 0.13 (θ=0.56) |
| H1N1 A/Boston/YGA-01050/2012 | (1.12, 0.60) | 0.33 (θ=2.31) | 0.16 (θ=0.74) |
| H1N1 A/Michigan/45/2015 | (1.07, 0.78) | 0.26 (θ=2.50) | 0.14 (θ=0.93) |
| H1N1 A/Idaho/07/2018 | (1.20, 0.16) | 0.37 (θ=2.16) | 0.18 (θ=0.59) |
| H3N2 A/Aichi/2/1968 | (-1.10, 0.89) | 0.20 (θ=0.33) | 0.13 (θ=1.90) |
| H3N2 A/Port Chalmers/1/1973 | (-1.04, 0.18) | 0.22 (θ=1.59) | 0.17 (θ=0.02) |
| H3N2 A/Bilthoven/1761/1976 | (-1.03, 0.05) | 0.16 (θ=1.54) | 0.14 (θ=3.11) |
| H3N2 A/Texas/1/1977 | (-1.04, 0.27) | 0.16 (θ=0.76) | 0.14 (θ=2.34) |
| H3N2 A/Netherlands/209/1980 | (-1.32, 0.25) | 0.22 (θ=1.30) | 0.19 (θ=2.87) |
| H3N2 A/Philippines/2/1982 | (-1.02, 0.21) | 0.23 (θ=1.57) | 0.17 (θ=0.00) |
| H3N2 A/Colorado/2/1986 | (-1.01, -0.39) | 0.17 (θ=1.72) | 0.13 (θ=0.14) |
| H3N2 A/Shanghai/11/1987 | (-1.06, 0.18) | 0.23 (θ=1.60) | 0.17 (θ=0.03) |

| | | | |
|---|---|---|---|
| H3N2 A/Sichuan/2/1987 | (-1.02, 0.05) | 0.24 (θ=1.63) | 0.16 (θ=0.06) |
| H3N2 A/Beijing/353/1989 | (-1.23, 0.29) | 0.22 (θ=1.34) | 0.18 (θ=2.91) |
| H3N2 A/Shangdong/9/1993 | (-1.16, 0.43) | 0.17 (θ=0.47) | 0.14 (θ=2.04) |
| H3N2 A/Johannesburg/33/1994 | (-1.21, 0.31) | 0.21 (θ=1.41) | 0.18 (θ=2.98) |
| H3N2 A/Brisbane/8/1996 | (-0.98, -0.33) | 0.16 (θ=1.65) | 0.12 (θ=0.08) |
| H3N2 A/Sydney/5/1997 | (-1.19, 0.78) | 0.23 (θ=0.56) | 0.17 (θ=2.13) |
| H3N2 A/Moscow/10/1999 | (-1.55, 0.82) | 0.26 (θ=0.55) | 0.18 (θ=2.12) |
| H3N2 A/Fujian/411/2002 | (-1.41, 1.46) | 0.28 (θ=0.30) | 0.13 (θ=1.87) |
| H3N2 A/California/07/2004 | (-1.22, 1.38) | 0.27 (θ=0.23) | 0.13 (θ=1.80) |
| H3N2 A/Wisconsin/67/2005 | (-1.08, 1.32) | 0.28 (θ=0.27) | 0.17 (θ=1.84) |
| H3N2 A/Brisbane/10/2007 | (-1.26, 1.07) | 0.27 (θ=0.37) | 0.18 (θ=1.94) |
| H3N2 A/Perth/16/2009 | (-1.21, 0.46) | 0.21 (θ=0.9) | 0.18 (θ=2.47) |
| H3N2 A/Indiana/10/2011 | (-0.82, 1.36) | 0.26 (θ=0.24) | 0.17 (θ=1.81) |
| H3N2 A/Victoria/361/2011 | (-1.21, 0.53) | 0.18 (θ=0.43) | 0.14 (θ=2.00) |
| H3N2 A/Texas/50/2012 | (-1.33, 1.44) | 0.28 (θ=0.27) | 0.13 (θ=1.84) |
| H3N2 A/Switzerland/9715293/2013 | (-1.16, 0.99) | 0.22 (θ=0.32) | 0.13 (θ=1.89) |
| H3N2 A/Hong Kong/4801/2014 | (-0.97, 0.39) | 0.16 (θ=0.75) | 0.14 (θ=2.32) |
| H3N2 A/Singapore/INFIMH-160019/2016 | (-0.99, 0.96) | 0.23 (θ=0.45) | 0.17 (θ=2.02) |
| H3N2 A/Perth/1008/2019 | (-1.34, 1.66) | 0.36 (θ=0.37) | 0.15 (θ=1.94) |
| Stem Ab 1 (CR8020) | (-2.26, -0.41) | 0.41 (θ=2.26) | 0.11 (θ=0.69) |
| Stem Ab 2 (315-27-1C08) | (-1.89, -0.47) | 0.24 (θ=2.36) | 0.10 (θ=0.79) |
| Stem Ab 3 (315-09-1B12) | (-1.68, -0.41) | 0.20 (θ=2.30) | 0.09 (θ=0.73) |
| Stem Ab 4 (315-19-1D12) | (-1.86, -0.97) | 0.31 (θ=2.60) | 0.11 (θ=1.03) |
| Stem Ab 5 (02-1D09) | (-1.65, -1.17) | 0.33 (θ=2.69) | 0.15 (θ=1.11) |
| Stem Ab 6 (04-1D10) | (-1.71, -1.35) | 0.45 (θ=2.76) | 0.15 (θ=1.19) |
| Stem Ab 7 (315-02-1F07) | (-1.49, -0.93) | 0.25 (θ=2.61) | 0.09 (θ=1.04) |
| Stem Ab 8 (15-5E04) | (-1.26, -0.92) | 0.23 (θ=2.54) | 0.12 (θ=0.97) |
| Stem Ab 9 (CT149) | (-1.24, -0.92) | 0.16 (θ=2.63) | 0.09 (θ=1.06) |
| Stem Ab 10 (13-1B02) | (-0.90, -0.84) | 0.19 (θ=2.63) | 0.12 (θ=1.06) |
| Stem Ab 11 (315-53-1A09) | (-0.90, -0.85) | 0.15 (θ=2.82) | 0.09 (θ=1.25) |
| Stem Ab 12 (21-1A10) | (-0.34, -0.60) | 0.15 (θ=2.38) | 0.13 (θ=0.81) |
| Stem Ab 13 (22-1B08) | (-1.08, -1.68) | 0.31 (θ=2.85) | 0.13 (θ=1.28) |
| Stem Ab 14 (54-4H03) | (-0.54, -1.36) | 0.20 (θ=2.85) | 0.12 (θ=1.28) |
| Stem Ab 15 (315-23-1C09) | (0.01, -1.24) | 0.14 (θ=0.07) | 0.09 (θ=1.64) |
| Stem Ab 16 (315-55-1E08) | (0.93, -1.37) | 0.17 (θ=0.28) | 0.10 (θ=1.85) |
| Stem Ab 17 (FI6v3) | (0.43, 0.31) | 0.16 (θ=1.57) | 0.08 (θ=3.14) |
| Stem Ab 18 (MEDI8852) | (0.26, -0.02) | 0.17 (θ=1.63) | 0.09 (θ=0.06) |

| | | | |
|---|---|---|---|
| Stem Ab 19 (315-53-1B06) | (0.31, -0.24) | 0.13 (θ=1.64) | 0.09 (θ=0.07) |
| Stem Ab 20 (315-53-1F12) | (0.46, -0.44) | 0.12 (θ=1.70) | 0.10 (θ=0.13) |
| Stem Ab 21 (315-55-1E11) | (0.65, -0.43) | 0.11 (θ=1.48) | 0.11 (θ=3.05) |
| Stem Ab 22 (CR9114) | (1.00, -0.25) | 0.12 (θ=0.58) | 0.10 (θ=2.15) |
| Stem Ab 23 (58-6F03) | (0.95, -0.55) | 0.17 (θ=0.15) | 0.13 (θ=1.73) |
| Stem Ab 24 (55-1D06) | (1.33, -0.64) | 0.22 (θ=0.42) | 0.12 (θ=1.99) |
| Stem Ab 25 (315-02-1H01) | (1.95, 0.17) | 0.21 (θ=1.26) | 0.09 (θ=2.83) |
| Stem Ab 26 (CR6261) | (2.12, -0.17) | 0.28 (θ=1.00) | 0.11 (θ=2.58) |
| Stem Ab 27 (02-1B02) | (2.07, -0.29) | 0.31 (θ=0.93) | 0.13 (θ=2.50) |
| Head Ab 1 (C05) | – | – | – |
| Head Ab 2 (CH65) | – | – | – |
| Head Ab 3 (F005-126) | – | – | – |
| Head Ab 4 (F045-092) | – | – | – |
| Head Ab 5 (310-33-1G04) | – | – | – |
| Head Ab 6 (5J8) | – | – | – |

**Dimensionality of the Landscape**

Since the Neutralization Landscape attempts to quantify the complex and potentially highly-multidimensional antibody-virus interactions in 2D, the position error could be greatly reduced in higher dimensional representations. To explore this possibility, we recreated the landscape from Figure 1C in 1D, 2D, 3D, and 4D (Figure S3). While a 1D map representation resulted in markedly higher error, the 2D, 3D, and 4D representations exhibited comparable error. Due to the ease of visualization, we opted to keep the 2D representation as is commonly done with antigenic cartography (*13*).

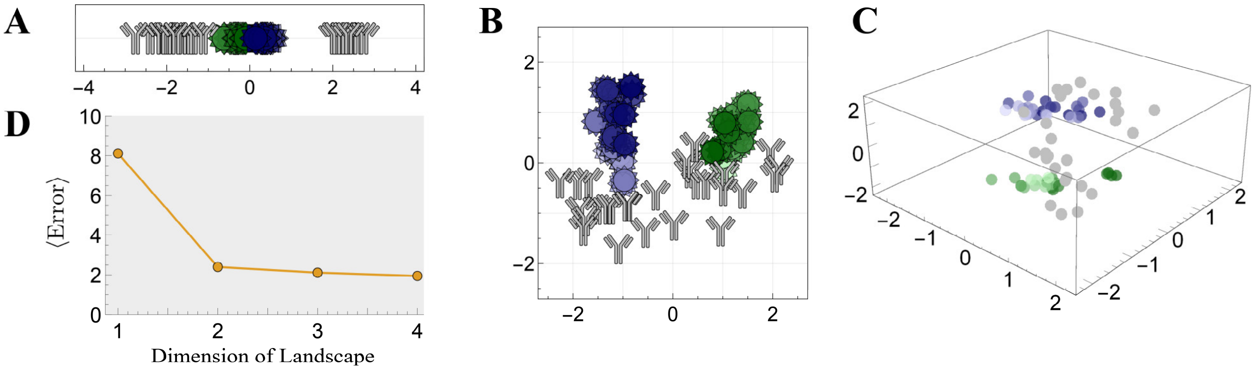


**Figure S3. Dimensionality of the Neutralization Landscape.** Monoclonal antibody neutralization data from Creanga *et al*. 2021 was combined with multidimensional scaling in (A) 1, (B) 2, or (C) 3 dimensions to create antigenic maps of the virus strains. In Panel C, the antibodies and viruses are all denoted by spheres. (D) The average fold-error of the landscape created in 1D-4D.

**Extrapolating the Behavior of New Antibodies**

Using the positions of the antibodies and viruses on the Neutralization Landscape, we can add additional entries using a few measurements to triangulate their coordinates (Figure 2). Triangulation works slightly better when the entries used for triangulation are spread apart on the map.

For example, since the H1N1 and H3N2 viruses lie on approximately vertical lines, we chose viruses spread out based on the *y*-coordinate of each virus. Thus, when choosing *N* viruses to triangulate a new antibody (with *N*=6 shown in the middle panel of Figure 2A), we chose *N*/2 H1N1 and *N*/2 H3N2 strains, with one virus chosen from the top third of viruses of each subtype, another chosen from the middle third, and the last chosen from the bottom third. Similarly, since the antibodies lie on an approximately horizontal line, we chose *N* antibodies spread out based on their *x*-coordinate when triangulating a new virus.

The triangulation and resulting predictions slowly improve in accuracy with the number of measurements used for triangulation. For example, a new virus triangulated using *N*=5 antibodies will have 78% of predictions with ≤4-fold error, which increases to 82% of predictions when *N*=10 antibodies are used (Figure S4).

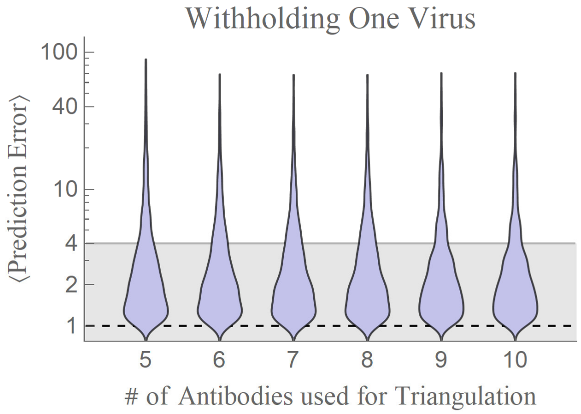


**Figure S4. Varying the number of triangulation measurements used to add a new antibody or virus.** We remove a virus from the dataset, triangulate its position using the number of measurements shown on the *x*-axis, and predict all other antibody-virus interactions. This process extends the antibody triangulation in Figure 2 (where we only considered *N*=6 measurements).

**The Triangle Inequality, Antibody-Virus Distance as a Metric**

For antibody-virus distance to form a metric, it must obey a triangle inequality. Colloquially, the triangle inequality ensures that the shortest distance between two points is a straight line, and that a more circuitous path does not represent a shortcut.

Given that antibody-virus distance corresponds to an experimentally measurable neutralization measurement (whereas antibody-antibody or virus-virus distance cannot be directly measured), we modify the usual triangle inequality to include two antibodies and two viruses (Figure S5, Equation 1). Although this relationship would more accurately be called a quadrilateral inequality, we will continue to refer to it as the more familiar triangle inequality. By analyzing all combinations of antibodies and viruses, we perform 400,000 tests of the triangle inequality, and we find that 98.8% satisfy Equation 1. If we account for the 2-fold error of the neutralization assay, the fraction increases to 99.7%, demonstrating that the triangle inequality is overwhelmingly satisfied on our antibody-virus data.

**Removing Head Antibody Neutralization**

Although virus-virus distance between viruses V1 and V2 cannot be directly measured, it nonetheless constrains the fold-difference in neutralization measurements (Figure 4B). For example, if two viruses lie near each other on the landscape ($d_{\text{V1-V2}}\approx 0$), then any stem antibody will have nearly identical distance to both viruses. Hence its neutralization should also be identical, resulting in a fold-difference≈1 between the antibody's $IC_{50}$s against both viruses (Figure S5B, where $d_{\text{V1-V2}}=0.4$). In the limit where $d_{\text{V1-V2}}=0$, we achieve a fold-difference=1 against all possible viruses.

As another example, if two viruses are 2 units apart, their $IC_{50}$s can differ by at most 100-fold (Figure S5C), with this maximum achieved when an antibody is collinear with the two viruses but does not lie between them (*e.g.*, brown antibody on the left). Antibodies lying on the perpendicular bisector to the segment connecting the two viruses will equally neutralize both viruses and hence achieve the minimum possible fold-difference of 1 (*e.g.*, pink antibody in the center).

When removing the neutralization of head antibodies, we use the constraint imposed by virus-virus distance. For example, if two viruses are $d_{\text{V1-V2}}=2$ units apart as in Figure S5C, and if a head+stem mixture exhibits an $\text{IC}_{50,\text{V1}}^{\text{mixture}} = 10^{-11}\text{M}$ and $\text{IC}_{50,\text{V2}}^{\text{mixture}} = 10^{-7}\text{M}$, then the stem antibody alone is bounded below by $\text{IC}_{50,\text{V1}}^{\text{stem Ab}} \geq \text{IC}_{50,\text{V2}}^{\text{mixture}} 10^{-d_{\text{V1-V2}}}\text{M} = 10^{-9}\text{M}$ (always using the larger $\text{IC}_{50}^{\text{mixture}}$ value), since the two measurements can be at most $10^{d_{\text{V1-V2}}}$ =100-fold different and both the stem and head antibodies weakly neutralize V2. However, this does not account for the noise in the assay, and hence we take the more conservative approach and only increase an $IC_{50}$ if it is more than $f_{\text{noise}}$=10-fold lower than this lower bound. In the example above, we would set $\text{IC}_{50,\text{V1}}^{\text{stem Ab}} \geq \frac{\text{IC}_{50,\text{V2}}^{\text{mixture}}}{f_{\text{noise}} 10^{d_{\text{V1-V2}}}\text{M}} = 10^{-10}\text{M}$ to account for noise.

Note that we always use the larger mixture $IC_{50}$ to bound the stem antibody's neutralization. If a head+stem mixture exhibits a large $IC_{50}$ against a virus, then both the head and stem antibodies must be weak against that virus. However, if this mixture exhibits a small $IC_{50}$, either the head antibody or stem antibody (or both) could potently neutralize the virus, and in the former case we should not use the mixture's $IC_{50}$ to alter any other values. Thus, we always use the larger mixture $IC_{50}$ values as "ground truth" to subtract the head antibody signal.

On the landscape, we visualize the new $IC_{50}$ bounded from below as a "zone of exclusion" represented by a gold disk, so that the stem antibody should lie as close to the red circles [signifying an $IC_{50}$ within the dynamic range of the assay] and outside all gold disks [representing $IC_{50}$s bounded from below] (Figure 4C). For example, when removing the head antibody signatures in Figure 4C, the red circles around the blue H3N2 viruses (left panel) expanded due to the constraint on virus-virus distance and became lower bounds (shown by gold circles in the right panel). The $IC_{50}$s against the green H1N1 viruses did not change because they all obeyed the constraints of the neutralization landscape. This (correctly) implies that the mixture's neutralization against these H1N1 viruses is dominated by the stem antibody, and hence there is no need to correct for the head antibody's neutralization.

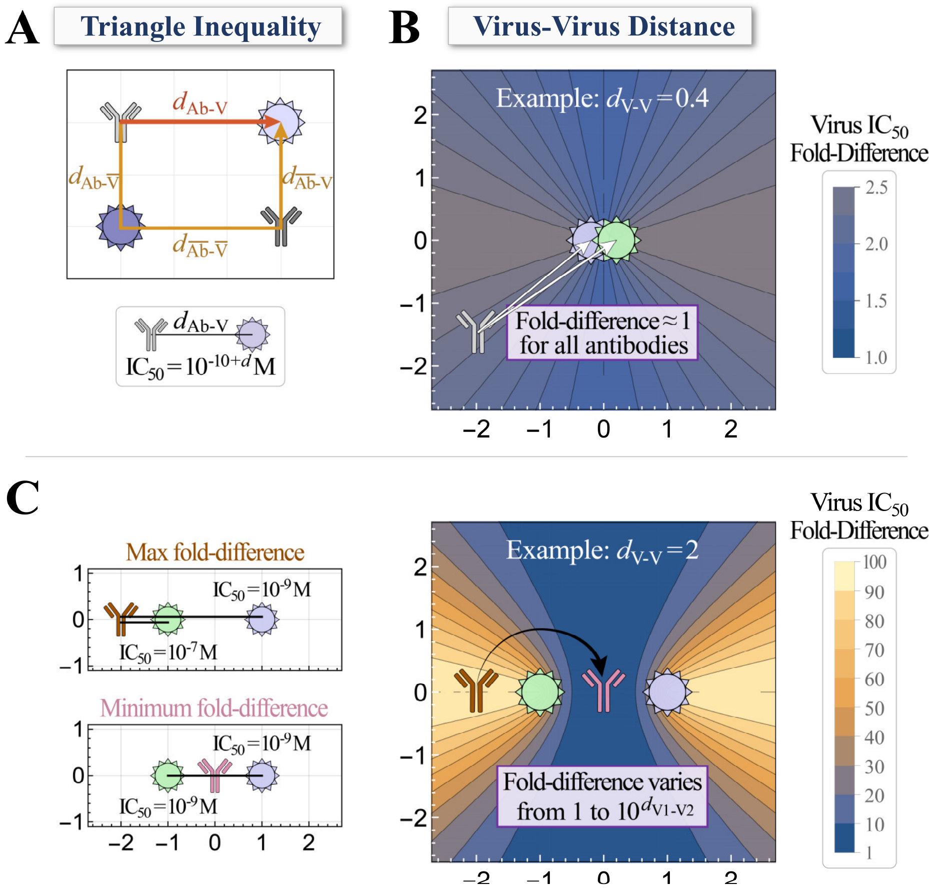


**Figure S5. Interpreting antibody-virus and virus-virus distance.** (A) The triangle inequality is defined between an antibody (Ab, top-left) and virus (V, top-right) relative to any other antibody ($\overline{\text{Ab}}$, bottom-right) and virus ($\overline{\text{V}}$, bottom-left). (B) Virus-virus distance constrains the fold-difference in neutralization ($IC_{50,\text{Virus 1}}/IC_{50,\text{Virus 2}}$). When two viruses lie at nearly the same coordinate, all stem antibodies will neutralize them identically. (C) More generally, when two viruses are separated by a distance $d_{V\text{-}V}$, the fold-difference in neutralization for these two viruses can vary from the maximum value $10^{d_{V\text{-}V}}$ (brown antibody) to the minimum value of 1 when both viruses are equally neutralized (pink antibody).

Finally, recalling that antibodies and viruses are treated symmetrically in our framework, we note that antibody-antibody distance has an analogous interpretation, constraining how differently any virus can be neutralized by both antibodies. Two antibodies at the same location will exhibit identical neutralization profiles, whereas two antibodies a distance $d_{\text{Ab-Ab}}$ apart will neutralize any virus at most $10^{d_{\text{Ab-Ab}}}$-fold differently.

**The Tradeoff between Antibody Potency and Breadth**

Using the Neutralization Landscape, we can computationally explore how potently any set of viruses can be neutralized by an optimal antibody (*i.e.*, an antibody with the smallest possible $IC_{50}$ against every virus in the set).

For example, in Figure 3 we explored how well an H3N2-specific antibody can inhibit all H3N2 vaccine strains from the 2004-05 to the 2018-19 season (A/Fujian/411/2002, A/California/07/2004, A/Wisconsin/67/2005, A/Brisbane/10/2007, A/Perth/16/2009, A/Victoria/361/2011, A/Texas/50/2012,

A/Switzerland/9715293/2013, A/Hong Kong/4801/2014, and A/Singapore/INFIMH-160019/2016). We similarly determined the optimal antibody for the H1N1 vaccine strains from this same period (A/New Caledonia/20/1999, A/Solomon Islands/03/2006, A/Brisbane/59/2007 [substituted with A/New York/08-1326/2008, see next paragraph], A/California/07/2009, and A/Michigan/45/2015).

Of these 15 vaccine strains, 14 were in our virus panel. The one missing strain, A/Brisbane/59/2007, was substituted with A/New York/08-1326/2008 whose HA sequence only differs by 2 amino acids. Given this sequence similarity, the two viruses should have similar neutralization profiles and hence have similar positions on the landscape. In general, the ability to substitute viruses based on their sequences greatly expands the utility of the Neutralization Landscape.

### B. Modeling Combinations of Antibodies

The workflow to decompose polyclonal serum can be split into three parts. First, we use monoclonal antibody data to create the Neutralization Landscape for the HA stem, and we use its structure to remove the neutralization from antibodies that do not target the HA stem (discussed in Section A above). Second, we determine how multiple stem antibodies collectively neutralize a virus (the purview of this section). Third, we combine the two previous points to determine which combination of stem antibodies best describes a mixture's neutralization data (explained in section C).

Here, we tackle the second part, namely, to create a model that takes the $IC_{50}$s of individual antibodies against a virus and predicts the $IC_{50}$ of their combination. As described below, we utilize measurements from our 2-antibody and 3-antibody mixtures to conclude that antibody combinations are well characterized by a competitive binding model, where all antibodies compete for the same region on HA.

**Overview of Antibody Mixtures**

We created 14 mixtures containing either two antibodies (at 1:1 stoichiometry) or three antibodies (at 1:1:1 stoichiometry) from our panel (Figure S6). We measured the neutralization of each combination against our virus panel as described previously (*9*). The $IC_{50}$ of these antibody mixtures together with the $IC_{50}$s of the monoclonal antibodies are provided in the Supporting Information file named "*(2) Creanga Neutralization Data - Monoclonal Antibodies and Mixtures.csv*".

Throughout this work, the mixture $IC_{50}$s represent the total concentration of IgGs at which 50% of the virus was neutralized. For example, Mixture #7 comprising Stem Ab 26 + Head Ab 1 neutralized H1N1 A/Weiss/1943 with an $IC_{50}$ of $5.25 \cdot 10^{-9}$ M, so that each individual antibody was present at half this concentration at the point of 50% neutralization.

The following section describes the competitive binding model we developed to test how each antibody's individual neutralization could predict the mixture's combined neutralization. Since all antibodies on our panel bind to the same region on the HA stem (*18–20*), we assumed competitive binding where only one antibody can bind an HA monomer at a time due to steric exclusion (Figure 5A) (*16*). We note that IgG is a bulky molecule, and the size of each of its two Fabs (8 nm×5 nm×4 nm) is roughly comparable to an HA trimer (cylinder with length 13.5 nm and diameter 5.5 nm) (*16*). Thus, there is little room for two antibodies to bind to the stem [Figure S8 of Ref. (*21*)], especially considering that HA trimers are tightly packed on the surface of the influenza virus with a mean separation of ~14 nm (*16*).

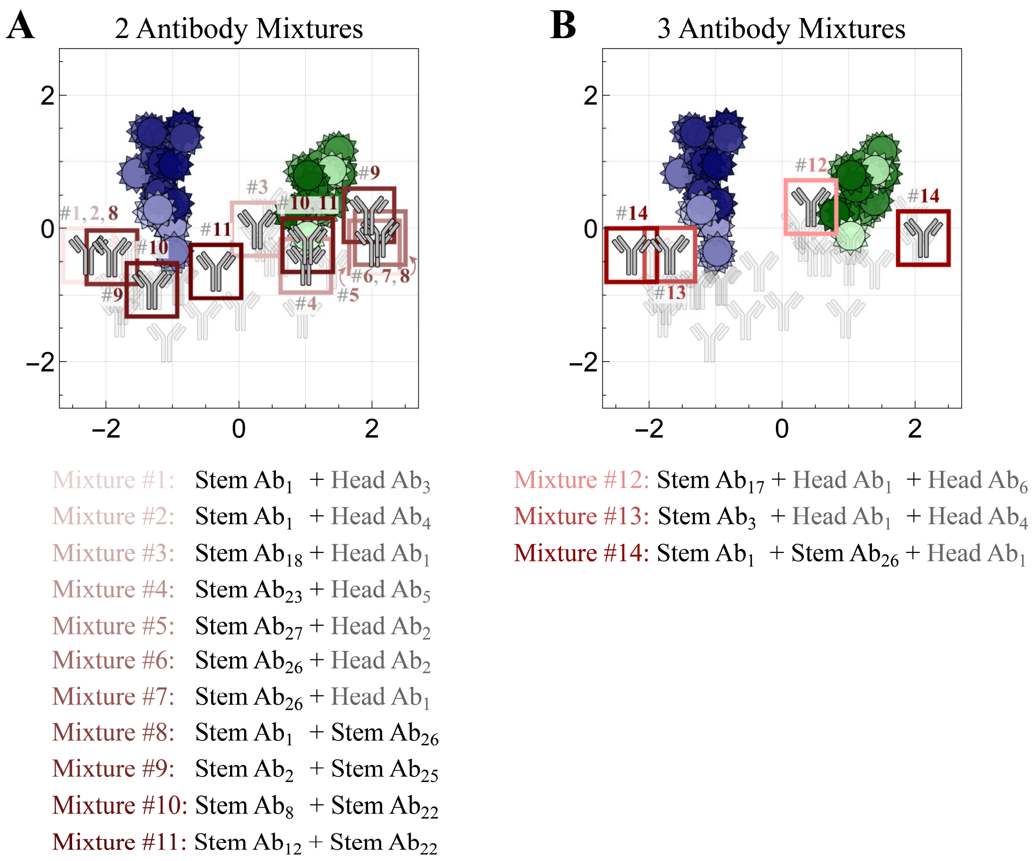


**Figure S6. Mixtures of two or three antibodies created to test the decomposition algorithm.** We created 14 mixtures of head and stem antibodies containing (A) 2 antibodies or (B) 3 antibodies. The stem antibodies used in these mixtures are boxed and labeled with the mixtures containing them. See Table S1 for the full antibody names.

**Collective Neutralization of Stem Antibody Mixtures is Well Characterized by Competitive Binding**

We begin by considering a two-antibody mixture with total IgG concentration $c$. This mixture is composed of a fraction $f_1$ of the first antibody and a fraction $f_2=1-f_1$ of the second antibody, so that the antibodies are at concentrations $c_1=f_1c$ and $c_2=f_2c$. Let $\mathrm{IC}_{50}^{(1)}$ be the concentration of the first antibody necessary for 50% inhibition (by itself) and $\mathrm{IC}_{50}^{(2)}$ be the analogous value for the second antibody.

Figure 5A enumerates the possible states within each model, using a simplified representation for a binding site on viral HA (purple) where each antibody can bind at a single site. A virus is assumed to be neutralized if any antibodies are bound to it. Hence, neutralization is given by

$$\mathrm{Neutralization}_{\mathrm{Competitive}} = \frac{\frac{c_1}{\mathrm{IC}_{50}^{(1)}} + \frac{c_2}{\mathrm{IC}_{50}^{(2)}}}{1 + \frac{c_1}{\mathrm{IC}_{50}^{(1)}} + \frac{c_2}{\mathrm{IC}_{50}^{(2)}}}$$

in the competitive binding model.

For each model, the mixture $\mathrm{IC}_{50}$ is defined as the total IgG concentration $c$ for which the neutralization formula equals 0.5, representing that a virion has a 50% chance of being neutralized. Hence

$$\mathrm{IC}_{50,\mathrm{Competitive}}^{\mathrm{Mixture}} = \left( \sum_j \frac{f_j}{\mathrm{IC}_{50}^{(j)}} \right)^{-1}$$

for the competitive binding model, and this formulation holds for mixtures with an arbitrary number of antibodies. With this equation, we predicted each mixture's $IC_{50}$ from the $IC_{50}$s of the constituent antibodies. For our stem+stem mixtures (#8-11 in Figure S6, $N$=165 data points), over 98% of measurements had ≤4-fold error [Figure 5B]. This tight agreement implies that if there are any errors in the monoclonal antibody or antibody mixture data, the competitive binding model can readily flag those outliers so they can be remeasured.

### C. Decomposing Defined Antibody Mixtures

In this section, we combined the Neutralization Landscape (described in section A) with the mathematical framework to model multiple stem antibodies (section B). With this basis set of antibody behaviors, the total neutralization of an arbitrary antibody mixture against a virus panel can be decomposed into the minimal set of antibodies that can give rise to the observed measurements.

**Why Polyclonal Mixtures need to be Represented using Multiple Points**

To date, antigenic cartography has been based on the interactions between polyclonal ferret sera and different influenza strains, with each virus and each serum represented by a single point on a map (*2*, *22*). This representation of polyclonal sera raises an important question: if two sera are pooled together, could any single point accurately represent their mixture?

As a simple example, consider the scenario shown in Figure S7 where serum 1 potently neutralizes virus 1 while serum 2 strongly neutralizes virus 2, with virus 3 lying somewhere in between. Traditional antigenic cartography would suggest that the combination of serum 1+2 would lie between the two points representing the original sera, but any such point (and, in fact, any single point on the map) fails to capture the expected strong neutralization against viruses 1 and 2 and the weak neutralization against virus 3.

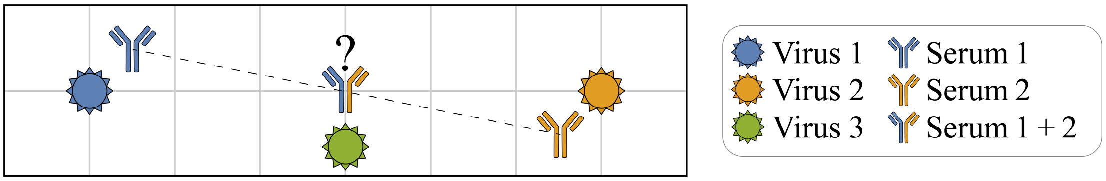


**Figure S7. Contradictions arise when representing polyclonal sera as single points.** Serum 1 is individually potent against virus 1, while serum 2 is individually potent against virus 2. Thus, combining both sera (1:1 stoichiometry) should result in a serum potent against both viruses. If we represent this combined serum as a single point, then minimizing the error to viruses 1 and 2 would imply that it lies between the two viruses [blue/yellow marker in the center]. This incorrectly implies that the combined serum will weakly neutralize viruses 1 and 2, and moreover it suggests that the combination will neutralize virus 3 much more potently than should be possible by either serum 1 or 2 alone. Instead, the correct representation of the combined serum should be to use two markers (at the locations of serum 1 and 2).

Extending this logic to its extreme, if all serum samples analyzed in Smith *et al*. 2004 were combined, they should effectively inhibit all H3N2 strains analyzed, yet no single point on their map could be sufficiently close to the spread-out viruses to match this expected level of protection.

Instead, the cartography approach based on monoclonal antibody data suggests that mixtures of antibodies should be represented by multiple points. For example, the mixture of serum 1+2 in Figure S7 should be represented by two markers at the locations of serum 1 and serum 2. The more polyclonal a mixture is, the more potential error may be introduced when representing it by a single point. The following sections examine the consequences of this multi-point representation for polyclonal mixtures.

**Defining Decomposition Error**

We first define the error metric used to quantify how our predicted antibodies compare to the true mixture behavior. We compute the mean fold-error between the neutralization of our predicted stem antibodies and the mixture's measured neutralization,

$$\langle \text{decomposition error} \rangle = 1/N \sum \text{fold-error}[\mathrm{IC}_{50}^{\text{predicted}}, \mathrm{IC}_{50}^{\text{measured}}].$$

Note that $\mathrm{IC}_{50}^{\text{measured}}$ quantifies the neutralization from just the stem antibodies within each mixture. This is experimentally accessible, since for each of our head+stem mixtures, we also created the stem mixture without the head component (Figure S6). More precisely, for mixtures #1-7 and #12-13 we know how the single stem antibody behaves from our monoclonal antibody data, while for mixture #14 we know how stem antibodies 1 + 26 behave from mixture #8. (The remaining mixtures #8-11 do not include any head antibodies.) Therefore, in each case, we can compare the predictions of the stem antibodies within a mixture against the measurements of a mixture containing only those stem antibodies.

$\mathrm{IC}_{50}^{\text{predicted}}$ incorporates all predicted stem antibodies and their stoichiometry via the competitive binding model (see the formula for $\mathrm{IC}_{50,\text{Competitive}}^{\text{Mixture}}$ in Section B). Following Smith *et al.*, whenever a measurement was given as a bound and the prediction obeyed this bound (*e.g.*, $\mathrm{IC}_{50}^{\text{measured}} > 10^{-7}$ M and $\mathrm{IC}_{50}^{\text{predicted}} = 10^{-6}$ M), we excluded measurement-prediction pairs from the summation above to prevent artificially *deflating* the error. However, when the measurement was given as a bound and the prediction did not satisfy this bound (*e.g.*, $\mathrm{IC}_{50}^{\text{measured}} > 10^{-7}$ M and $\mathrm{IC}_{50}^{\text{predicted}} = 10^{-8}$ M), we included this pair in the error calculation with $\mathrm{IC}_{50}^{\text{measured}}$ equal to its bound.

**Overview of the Decomposition Algorithm**

Serum decomposition proceeds by characterizing a mixture using an increasing number of antibodies, halting once the addition of another antibody no longer markedly decreases the decomposition error.

1. We first remove any head antibody neutralization (described in SI Section A) and then determine the initial number of antibodies for decomposition. Ordinarily we start by describing a mixture with $n=1$ antibody and then consider $n=2, 3\ldots$ antibodies (as shown in Figure S8A,B). However, if H1N1 virus neutralization indicates a stem antibody on the right half of the landscape while H3N2 virus neutralization points to an additional antibody on the left half of the landscape, we begin with $n=2$ antibodies and then consider $n=3, 4...$ antibodies. This process is shown by the 3D panels on the left of Figure S8, and is discussed in more detail below.
2. Starting from $n$ antibodies, we accept the best decomposition with $n+1$ antibodies if the ⟨decomposition error⟩ decreases by at least 20%. This ensures that adding another antibody markedly reduces the error.

When the decomposition with $n+1$ antibodies decreases ⟨decomposition error⟩ by less than 20%, the search terminates and returns the best decomposition containing $n$ antibodies.

At each step, we consider all possible positions for the $n$ antibodies and (for $n \geq 2$) all possible stoichiometries, but we force each antibody to comprise ≥10% of the mixture to ensure that its neutralization can be clearly detected (since an antibody comprising ~1% of a mixture will negligibly affects the mixture's neutralization). For example, for a mixture comprising 1 head+1 stem antibody, we consider everything from a 10%/90% composition to a 90%/10% composition.

Figure S8 shows three example decompositions for mixture #6 [top row], mixture #14 [middle row], and mixture #8 [bottom row]. In the top row, we remove the head neutralization signal and fit a plane through the H1N1 $IC_{50}$s and another plane through the H3N2 $IC_{50}$s (as shown on the 3D panel on the left). Because the dynamic range of our assay cannot detect weak $IC_{50}$s ($>1.6 \cdot 10^{-7}$ M), and because including these weak values would skew the plane to be artificially horizontal, we remove these large $IC_{50}$s and only fit a plane if there are ≥5 values remaining. For mixture #6, we cannot fit a plane through the H3N2 viruses, and hence we begin with a decomposition using $n=1$ antibody. The transition from $n=1\to2$ antibodies decreases the error by only 5%, correctly indicating that the mixture contains 1 stem antibody (shown in gray).

For mixture #14 shown in the middle row of Figure S8, the H1N1 and H3N2 planes both tilt to the right, and hence we begin with $n=1$ antibody. The transition from $n=1\to2$ decreases error by 25% and is accepted, but $n=2\to3$ only decreases the error by an additional 10% and is rejected. Thus, the mixture is correctly predicted to have 2 stem antibodies, although the positions of these antibodies is markedly off. The fact that both the red and gray antibodies have nearly identical neutralization profiles (as seen by the small decomposition error) demonstrates that this set of neutralization measurements can arise from multiple sets of antibodies.

Mixture #8, shown in the final row of Figure S8, has an H1N1 plane tilting to the right (the normal vector to the plane with a positive $z$-component also has a positive $x$-component) whereas the H3N2 plane tilts to the left (the normal vector to the plane with a positive $z$-component has a negative $x$-component), suggesting that there is one antibody on the left half of the landscape and another on the right half. Hence, we begin with $n=2$ antibodies. The transition $n=2\to3$ does not decrease the error by 20% and is rejected, so that the mixture is correctly predicted to have 2 antibodies.

We note that while we have only quantified the optimal decomposition, we can also quantify how good the "next best" alternative is (containing either a different number of stem antibodies or a different position/stoichiometry for these antibodies) by comparing their ⟨decomposition error⟩. Thus, we can clarify when the transition between $n$ and $n+1$ antibodies is just shy of the 20% threshold, or if different configurations of $n$ antibodies have nearly identical error, thereby quantifying the degeneracy of the antibody response.

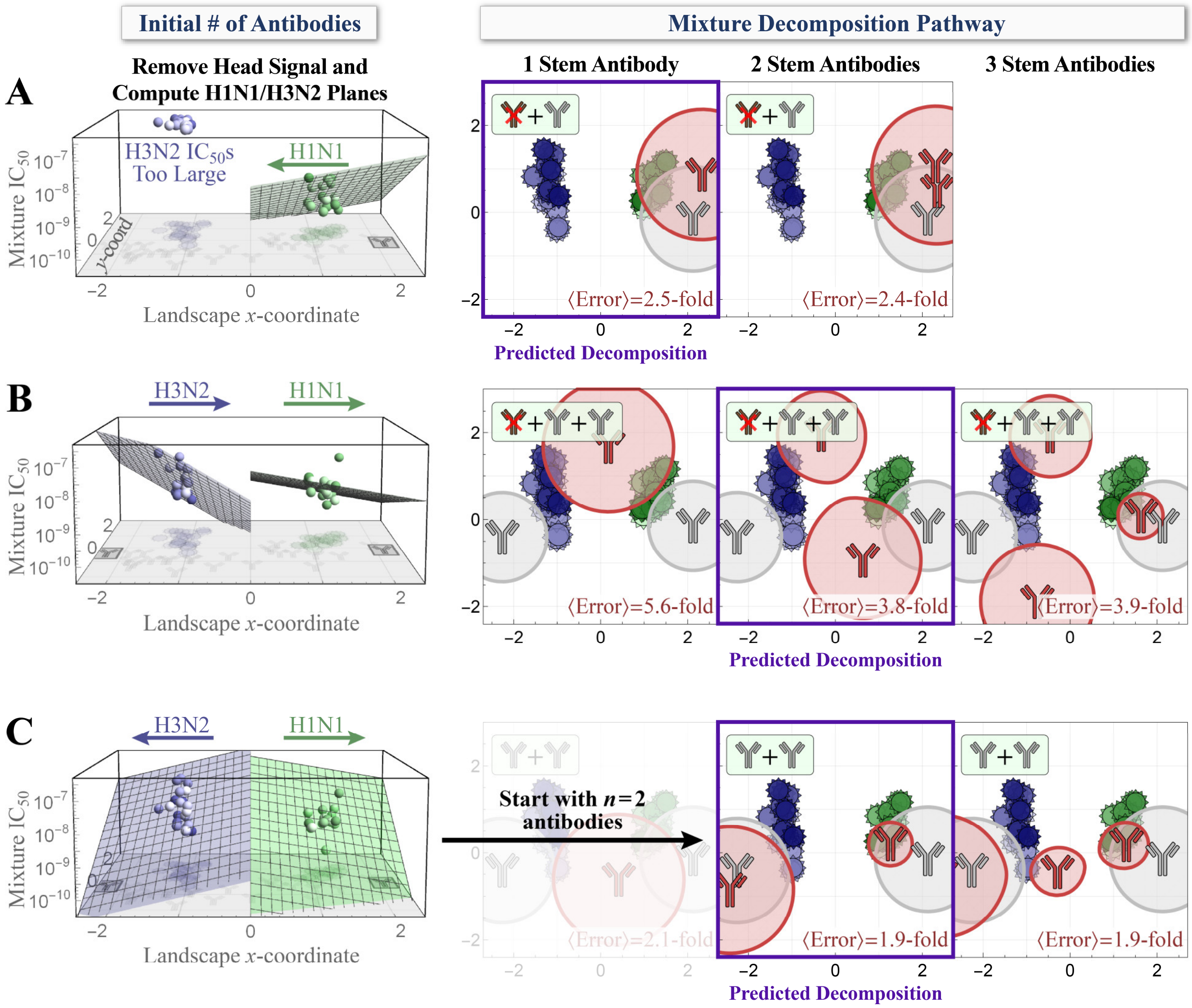


**Figure S8. Decomposition algorithm to detect the number of stem antibodies in a mixture.** We decompose the collective neutralization of a mixture against our virus panel using an increasing number of antibodies, stopping when the error will not decrease by at least 20% with the addition of the next antibody. Examples are shown for (A) mixture #6, (B) mixture #14, and (C) mixture #8. *Left*, By default, we begin by using a decomposition with $n$=1 antibody (Panels A,B) unless a plane fit through the H1N1 $IC_{50}$s tilts towards the right of the landscape while a plane through the H3N2 $IC_{50}$s tilts towards the left (as in Panel C), in which case we begin with $n$=2 antibodies. On each 3D plot, the 2D map is shown on the $x$-$y$ plane, and the stem antibodies in the mixture are boxed in gray. *Right*, Decomposition proceeds by analyzing $n$, $n$+1, $n$+2… antibodies, halting once the decomposition error decreases by less than 20%. The predicted decomposition is boxed in purple. On each 2D landscape, the stem antibodies in the mixture are shown in gray while the antibodies predicted by decomposition are shown in red. The circles around each antibody show the region of virus neutralization when the mixture has a total concentration of $10^{-8.5}$ M.

### Results from Decomposing Antibody Mixtures

Figure S9 shows the decompositions for our 14 antibody mixtures. On each plot, we show the location of the actual (gray) and inferred (red) antibodies that collectively give rise to the mixture's neutralization profile. Panel D shows the number of inferred antibodies and their stoichiometry. These decomposition results are also available in the Supporting Information file named "*(3) Decomposition Results.csv*", where each antibody in the decomposition is classified using the format {(map coordinates), fractional composition of the stem response}.

For example, as shown for mixture #14 in Figure S9C, we predict that there are two stem antibodies with 70%/30% stoichiometry, whereas the true mixture has an equal 50%/50% composition (since this head+stem+stem mixture has equal amounts of each antibody, so after removing the head antibody the two stem antibodies comprise 50%/50% of the mixture). The radius of the circular regions surrounding each antibody corresponds to the region of ≥50% neutralization when the total IgG concentration of stem antibodies in the mixture equals $10^{-8.5}$ M. Thus, the gray circles surrounding the actual antibodies have a radius of $1.5+\log_{10}(½)=1.2$, where 1.5 corresponds to the total mixture concentration of $10^{-8.5}$ M while $\log_{10}(½)$ accounts for the fact that each antibody only comprises half the mixture. The radius of the red circles around the predicted antibodies will be $1.5+\log_{10}(3/10)=1.0$ and $1.5+\log_{10}(7/10)=1.3$ for the antibodies on the top and bottom, respectively. In addition, the two red circles are slightly elongated towards each other because the other antibody can stretch the region of ≥50% virus neutralization, as seen by the solid versus dashed lines in Figure 5C. (The gray circles are minimally elongated because those antibodies are further apart. This is similar to the circles in Figure 5C when the total antibody concentration equals $10^{-9.5}$ M, where the dashed and solid lines overlap.) For each decomposition, we only show the viruses that a mixture was measured against — mixtures #4, 5, 10, and 11 were measured against 24 viruses, while the other 10 mixtures were measured against 39 viruses.

In a single case of a mixture with multiple antibodies (mixture #9), we predicted the wrong number of stem antibodies, inferring one instead of two stem antibodies. Moreover, if we decompose the neutralization of the 27 monoclonal antibodies in our panel, 22/27 are correctly decomposed as containing a single stem antibody while the other 5 are overfit as having two stem antibodies. Such cases happen when the 2D map representation for an antibody has large error, or when a different antibody configuration happens to more closely match the antibody's neutralization profile. In the majority of cases, decomposition determines the correct number of stem antibodies, albeit with large deviations in the stoichiometry. Notably, in *all* cases the resulting decompositions have ≲5-fold error compared to the true mixture. Given that the collective neutralization measurements must determine the number of antibodies, their locations on the map, and their fractional compositions, this low error highlights the wealth of information encoded by the virus coordinates on a Neutralization Map.

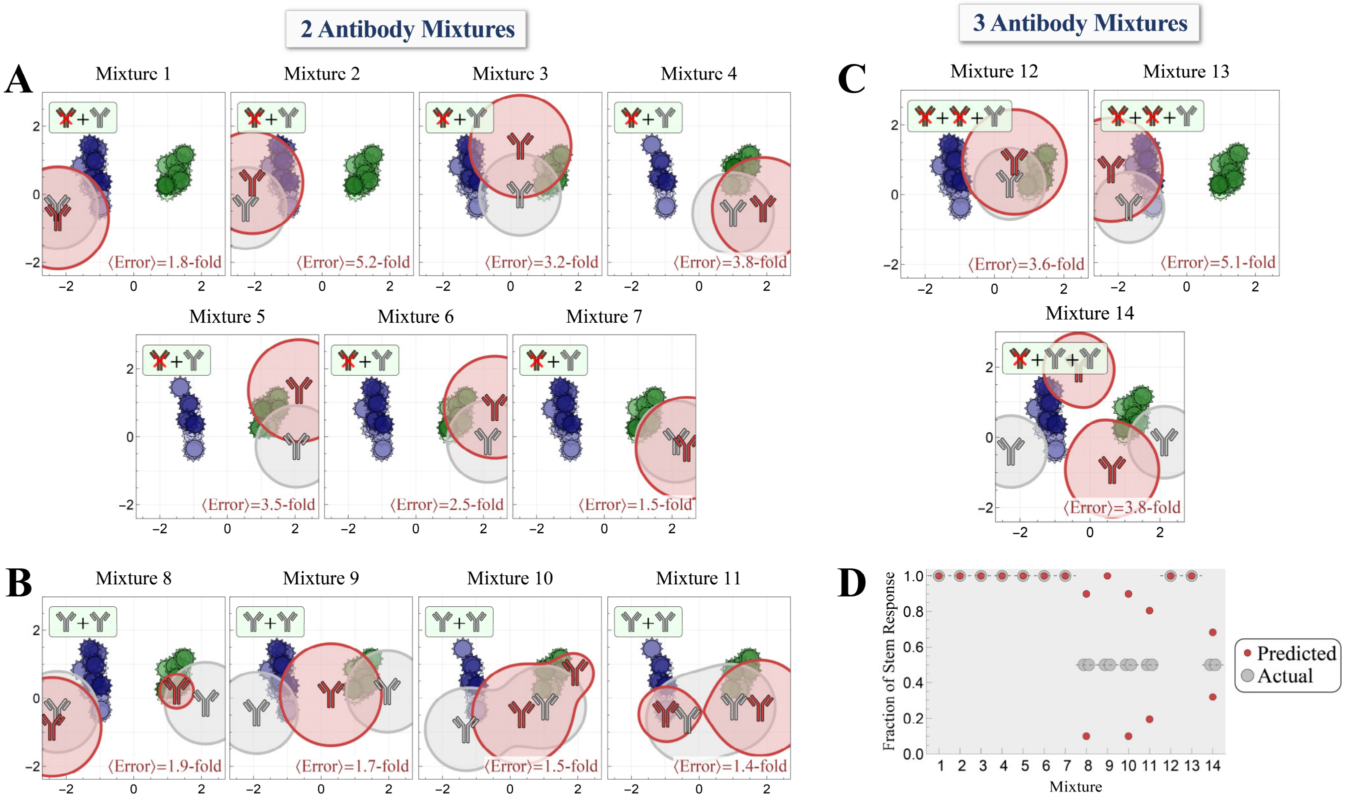


**Figure S9. Decomposing of all antibody mixtures.** For each mixture, we computationally remove the neutralization from any head antibodies and predict the optimal number, stoichiometry, and neutralization profiles of the stem antibodies within. Decomposition was performed without any knowledge of the antibodies within the mixture. Results for (A) head+stem, (B) stem+stem, or (C) head+head+stem or head+stem+stem mixtures. (D) The resulting stoichiometry of the predicted vs actual antibodies, shown as the fraction of the stem antibody response (since the neutralization from head antibodies was removed).
